# Spatial multi-omic map of human myocardial infarction

**DOI:** 10.1101/2020.12.08.411686

**Authors:** Christoph Kuppe, Ricardo O. Ramirez Flores, Zhijian Li, Monica Hannani, Jovan Tanevski, Maurice Halder, Mingbo Cheng, Susanne Ziegler, Xiaoting Zhang, Fabian Preisker, Nadine Kaesler, Yaoxian Xu, Remco M. Hoogenboezem, Eric M.J. Bindels, Rebekka K. Schneider, Hendrik Milting, Ivan G. Costa, Julio Saez-Rodriguez, Rafael Kramann

## Abstract

Myocardial infarction is a leading cause of mortality. While advances in the acute treatment have been made, the late-stage mortality is still high, driven by an incomplete understanding of cardiac remodeling processes^1,2^. Here we used single-cell gene expression, chromatin accessibility and spatial transcriptomic profiling of different physiological zones and timepoints of human myocardial infarction and human control myocardium to generate an integrative high-resolution map of cardiac remodeling. This approach allowed us to increase spatial resolution of cell-type composition and provide spatially resolved insights into the cardiac transcriptome and epigenome with identification of distinct cellular zones of injury, repair and remodeling. We here identified and validated mechanisms of fibroblast to myofibroblast differentiation that drive cardiac fibrosis. Our study provides an integrative molecular map of human myocardial infarction and represents a reference to advance mechanistic and therapeutic studies of cardiac disease.

---

Coronary heart disease driving acute myocardial infarction (MI) is the largest contributor to cardiovascular mortality, which in turn is the leading cause of all deaths worldwide.^3,4^ Tremendous progress has been made in the acute therapy of MI focusing primarily on percutaneous coronary intervention and resulting in a decreased acute mortality.^5^ However, the morbidity and mortality caused by left ventricular cardiac remodeling post-MI remain unacceptably elevated.^6,7^ Cardiac remodeling after MI involves immune cell recruitment and demarcation of the infarcted area followed by tissue digestion, phagocytosis, myofibroblast activation, scar formation and neovascularization.^8^ Understanding the exact cellular and molecular mechanisms of cardiac remodelling processes from the acute ischemic event to the chronic cardiac scar formation in their spatial context will be key to developing novel therapeutics.

Here we used a combination of single-cell gene expression and chromatin accessibility technologies and spatially resolved transcriptomics to study the cell-type specific changes in gene regulation, providing an integrated map of cardiac remodeling after MI. We defined candidate cis-regulatory DNA elements (CREs, regions of non-coding DNA that regulate transcription of neighbouring genes) which revealed gene regulatory networks controlling specific cardiac cell types in health and disease. We projected this information onto specific tissue locations, thus allowing us to spatially map putative enhancers controlling gene regulation e.g. in the myocardial border zone area. This, in turn, enabled us to gain novel insights into gene-regulatory programs driving fibroblast to myofibroblast differentiation in cardiac scar formation. Our results provide a comprehensive spatially resolved characterization of gene regulation of the human heart in homeostasis and after myocardial infarction. We anticipate that this data will be a reference map for the field and can be utilized for future studies, and ultimately for the development of novel therapeutics.

## Results

### Integrative spatial and single-cell genomics of the human heart

We applied an integrative single-cell genomic strategy with single nuclear RNA sequencing (snRNA-seq) and single nuclear Assay for Transposase-Accessible Chromatin sequencing (snATAC-seq) together with spatial transcriptomics from the same tissue (10x Genomics Visium) to map human cardiac cells in homeostasis and after MI at unprecedented spatial and molecular resolution (Fig. 1a-c). We compared a non-transplanted donor heart (control, C) as control to the necrotic core (ischemic zone, IZ), border zone (BZ) and the non-affected left ventricular myocardium (remote zone, RZ) of patients with acute MI (Fig. 1a). These acute MI specimens were obtained from hearts explanted 2-5 days after the onset of clinical symptoms (chest pain), before the patients received total artificial hearts due to cardiogenic shock and as a bridge to transplantation. We also analyzed two human heart specimens at later stages after MI (3 months and 12 years, fibrotic zone FZ) with ischemic heart failure that were available due to orthotopic heart transplantation. A non-transplanted donor heart served as a control (clinical characteristics are outlined in Extended Data Fig. 1a). We obtained 10μm cryo-sections of each cardiac specimen and isolated nuclei from the remaining specimen directly adjacent to the cryo-section with subsequent fluorescent activated nuclei sorting (FANS) for snRNA- and snATAC-seq (Fig. 1b). We obtained gene expression data from 40,530 nuclei in total, with an average of 1,335 genes per nucleus together with open chromatin in overall 18,213 nuclei with an average of 24,333 fragments per nucleus (*n* = 7). The spatial transcriptomics datasets contained an average of 3874.5 spots per specimen and 1,813 genes per spot with a large variability mainly due to the underlying biological process with cell-death after MI (Extended Data Fig. 1b-e). We used an integrative data analysis approach spanning all three modalities of our single-cell experiments to study cardiac cell specific information and their interactions in their spatial context (Fig. 1c, Extended Data Fig. 2).

**Fig. 1.**
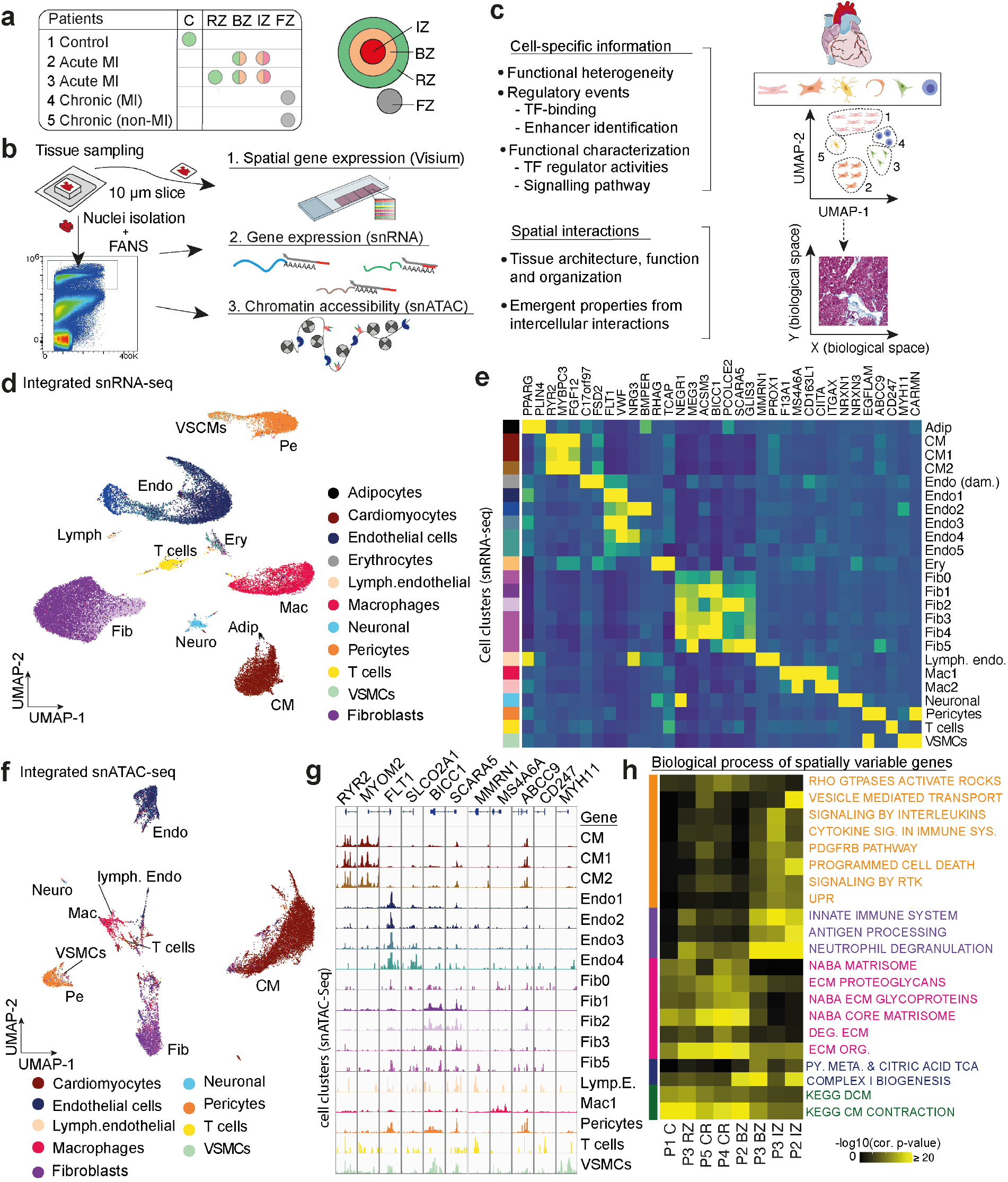
Multi-omic analysis of the human heart in health and disease. **a.** Overview of patients and samples in this study. In total 8 tissue samples were studied from 5 human hearts in three different modalities (snRNA-seq, snATAC-seq and spatial gene expression (Visium)). Areas included control (C), remote zone (RZ), border zone (BZ), ischemic zone (IZ) and fibrotic zones (FZ). **b.** A cryo-section of the selected tissue sample was used for spatial gene expression assay (Visium) and the contiguous remnant tissue was used for nuclei isolation followed by FANS (fluorescent activated nuclei sorting). **c.** Schematic of the integrative data analysis pipeline (for details see Figure S1). **d.** UMAP embedding of integrated 40,530 single nuclei transcriptomes from all 8 human heart tissue samples. Colors refer to annotated celltypes. CM (cardiomyocytes), Fib (fibroblasts), Neuro (neuronal cells), VSMCs (vascular smooth muscle cells), Mac (Macrophages), Pe (pericytes), T (T-cells), Endo (endothelial cells). **e.** Heatmap showing average expression of marker genes in each cell-type. **f.** UMAP embedding of 18,213 single nuclei chromatin accessibility profiles. **g.** Pseudo-bulk ATAC-seq tracks showing chromatin accessibility of selective marker genes. **h.** Heatmap showing overrepresented gene-sets in the spatially variable genes of each spatial gene expression dataset using hypergeometric tests. RTK = receptor tyrosine kinase, UPR = unfolded protein response, ECM = extracellular matrix, DEG = degradation, ORG = organization, PY = pyruvate, DCM = dilatative cardiomyopathy. Colours indicate biological processes; cell signalling process (orange), immune process (violet), cellular matrix process (pink), metabolic process (blue), and cardiacmuscle process (green).

### Single-cell transcriptome and chromatin landscape revealed heterogeneity of human cardiac cell-types

To establish a consistent map of cell types with snRNA-seq across samples, we first clustered the data for each sample individually and annotated the clusters with curated marker genes from the literature.^9–11^ To unify cell type annotations between samples, we integrated and batch-corrected data from all samples and clustered them based on the correlation of average gene expression (Fig. 1d, Extended Data Fig. 3a-b). This cluster analysis revealed 10 major cell types with several subpopulations (total *n* = 24 clusters) (Fig. 1d-e) which is largely in line with the recent literature on healthy human hearts.^9,11^ We next clustered the snATAC-seq data for each sample (Fig. 1f). snATAC clusters were annotated using the most representative labels following label transfer from the snRNA-seq data, allowing the identification of 8 of the 10 major cell types across samples identified in snRNA-seq data (Fig. 1f). The promoter accessibility of marker genes in pseudo-bulk ATAC-seq data confirmed the cell type identities (Fig. 1g). We observed a consistent clustering of cell types after combination and batch-correction of all snATAC-seq samples and identified cell-type specific transcription factor binding activities with HINT-ATAC^12^(Extended Data Fig. 3b-c). Of note, while major cell types were present across most specimens, various cell types were only identified in individual specimens, likely reflecting a distinct cell-state during injury or repair (Extended Data Fig. 3d). Together, our integrative single-cell analysis defined a consistent and non-redundant catalog of cell types that comprise the adult human heart across multiple modalities and samples.

We next investigated whether our spatial transcriptomics datasets reflected known biological processes of human myocardial infarction. To this end, we identified spatially variable gene expression across samples with SPARK^13^ and identified overrepresented biological processes using hypergeometric tests. This analysis revealed functional and organizational differences consistent with the underlying biological conditions (Fig. 1h). In the acute MI ischemic zone, we observed an enrichment of spatially variable gene expression associated with the innate immune system, neutrophil degranulation and programmed cell-death, and a depletion of fibrotic and muscle contraction processes (Fig. 1h). Consistently, the chronic remodeled heart (late stage after MI) showed enrichment of spatially variable genes associated with extracellular matrix (ECM) proteoglycans, glycoproteins and other matrisome components in line with the expected fibrotic processes captured in these specimens. The borderzone specimens showed an enrichment of genes associated with mitochondrial complex I biogenesis and pyruvate metabolism/citric acid TCA cycle, both confirming the response to injury and potentially altered redox state and metabolism of this area. In the control and remote zone specimens, we observed an enrichment of spatially variable genes associated with muscle contraction linked to an overrepresentation of healthy cardiomyocytes in these samples. Overall, this analysis confirms that the spatial data clearly reflects known zones of biological processes after myocardial infarction.

### Integrative multi-omic analysis of the healthy human heart

We first aimed to deeply characterize the integrated data of the control human heart specimens as a reference dataset (Extended Data Fig. 1a). We identified 12 cell types from snRNA-seq (*n* = 8,335) and 8 cell types from snATAC-seq (*n* = 3,849), respectively (Fig. 2a-b, Extended Data Fig. 4a). Transcription factor (TF) footprinting analysis of snATAC-seq data revealed TF binding activity of known cell-specific TFs such as MEF2C in cardiomyocytes, ETV6 in macrophages and SOX8 in endothelial cells (Fig. 2c, Extended Data Fig. 4b-c). The activity of these TFs was further confirmed by cell-type specific expression of their target genes (Fig. 2d, Extended Data Fig. 4c). Clustering of the spatial transcriptomic data of the same left ventricular heart tissue resulted in 11 molecularly distinct clusters (Fig. 2e). Differential gene expression analysis revealed that cardiomyocytes were the major transcriptomic contributors to individual spatial clusters (Fig. 2e, Extended Data Fig. 4d). Of note, marker genes allowed for the identification of common cell types in the distinct areas, like cardiomyocytes, but also rare cardiac cell populations such as vascular smooth muscle cells (<1% of cells in the snRNA-seq data). Interestingly, mast cells (*CPA3+*) were identified in a distinct spatial cluster (cluster 9, Fig. 2e) while they were not detectable in either the snRNA- or the snATAC-seq datasets (Fig. 2a-b). Comparison of the top differentially expressed genes in both the snRNA-seq and spatial transcriptomic dataset confirmed the spatial annotation (Fig. 2e, Extended Data Fig. 4d-e).

**Fig. 2.**
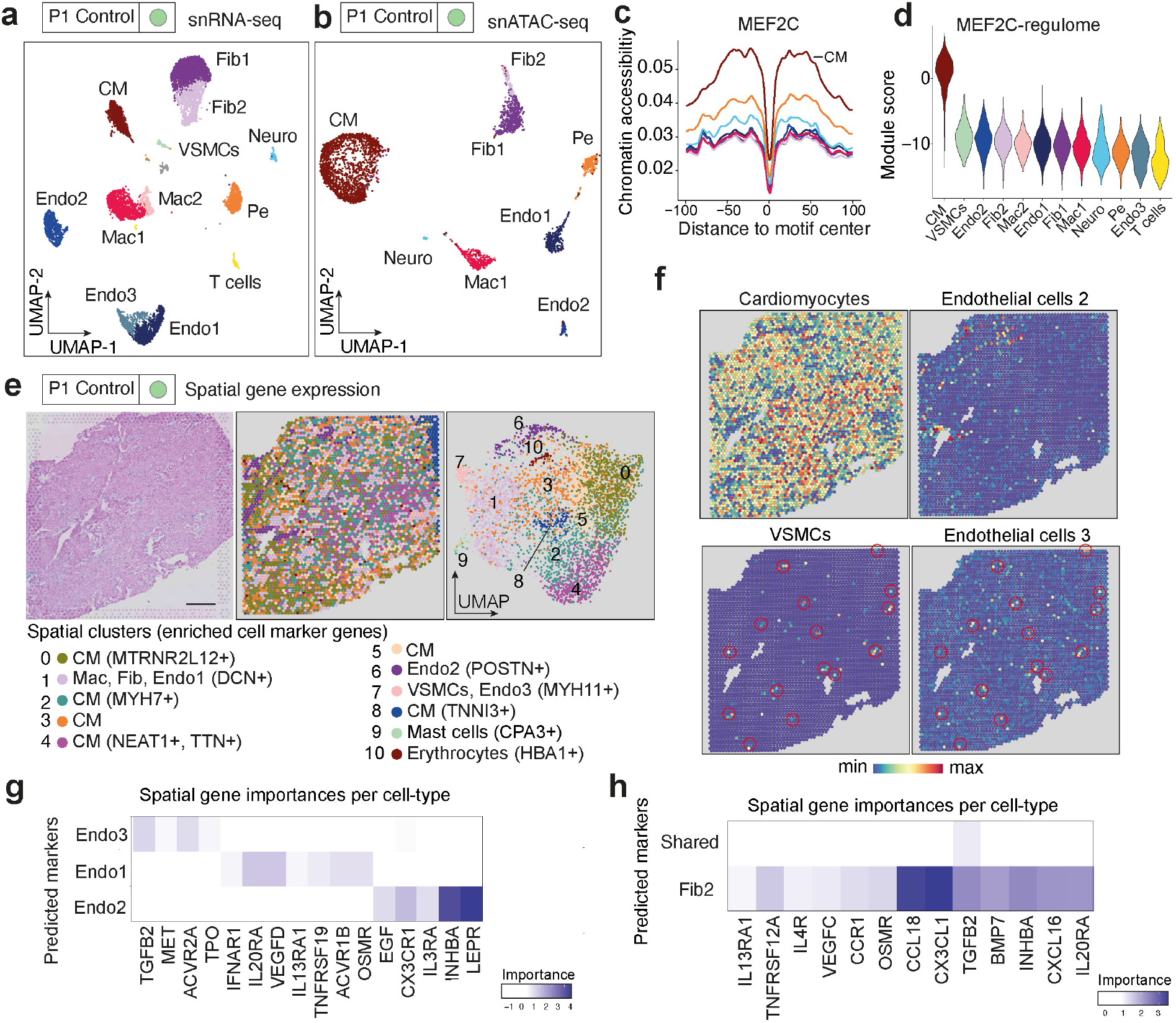
Multi-omic analysis of the healthy human heart. **a.** UMAP embedding of 8,335 single nuclei transcriptomes from a control human heart resulting in 12 clusters. Colors refer to annotated cell types. CM (cardiomyocytes), Fib (fibroblasts), Neuro (neuronal cells), VSMCs (vascular smooth muscle cells), Mac (Macrophages), Pe (pericytes), Endo (endothelial cells). **b.** UMAP embedding of 3,849 single nuclei open chromatin profiles resulting in 8 clusters. **c.** Line plot showing footprint of MEF2C from the control human heart. Y-axis represents the average ATAC-seq signal around the transcription factor binding sites. **d.** Violin plot showing summarized levels of expression of MEF2C-regulated genes per cell-type using module scores. **e.** HE-staining of the control heart tissue section (left). Clustering of each individual spatial detection spot resulted in 10 clusters (middle). Colors indicate the cluster assignments. UMAP embedding of spatial detection spots (right). Scale bar = 1mm. **f.** Cell-type score per spot by label transfer showing the distribution of cardiomyocytes, endothelial cells, VSMCs and endothelial cells 3. See Extended Data Fig. 5c for other cell-types. **g.** MISTy mean importances of the spatial expression of extracellular matrix and cytokine genes to predict the expression of marker genes of different subtypes of endothelial cells and **h.** fibroblasts.

We then integrated snRNA-seq and snATAC-seq data to generate a reference dataset which was used to map cell types to corresponding spatial transcriptomic data (Extended Data Fig. 5a-b). Since each individual spot on the spatial transcriptomic slide putatively contains multiple cells, we transferred the labels from the integrated data to spatial gene expression data and mapped different cell-types on each individual spatial location (Extended Data Fig. 5c). Interestingly, we observed three different populations of endothelial cells (endothelial cells 1-3; Extended Data Fig. 5c), in three distinct areas (spatial clusters 1, 6 and 7; Fig. 2e). Endothelial cells 3 were located in spatial cluster 7 (Fig. 2e), which was enriched with other cell types, yet showed an increased pathway activity of TGFβ signaling (Extended Data Fig. 5d), increased regulome-based activity of SMAD3 (Extended Data Fig. 5e) and close spatial colocalization with VSMCs (Fig. 2f, Extended Data Fig. 5c). This suggested a spatial arterial signaling niche and that endothelial cells 3 are arterial/arteriolar endothelial cells. We confirmed this finding by staining SEMA3G, an endothelial cells 3 specific marker, that indeed stained endothelial cells surrounded by ACTA2 positive VSMCs indicating an arterial location (Extended Data Fig. 5f). Of note, Litvinukova et al. also reported the presence of SEMA3G+ arterial endothelial cells recently^9^.

Endothelial cells 2 cells were defined by high periostin (*POSTN*) expression, suggesting a mesenchymal phenotype^14^ (Extended Data Fig. 4e), and expression of genes that might indicate an endocardial origin (*NPR3* and *CDH11*)^15^ (Extended Data Fig. 5g).

We next estimated to what extent the spatial neighborhood of specific cell-types in the tissue influenced their gene expression, using MISTy^16^. MISTy models the expression of cell-type markers using spatially contextualized views, for example in the detection spot itself (intraview) or the surrounding (paraview). In this model, an increment in the prediction of gene expression after using spatially contextualized views points at a potential relationship between a spot and its neighbors. This relationship may represent coordinated functions in areas of the tissue or intercellular interactions. After modeling the spatially resolved expression of endothelial cell marker genes, we observed that the local expression of *NPPA, LEPR*, and *INHBA* predicted the expression of endothelial cells 2 marker genes (Fig. 2g, Extended Data Fig. 5h). *NPPA* has been previously described as a marker of trabecular myocardium in the subendocardial zone^17^ thus suggesting an endocardial location of endothelial cells 2. To validate this hypothesis, we performed multiplex in situ hybridizations (ISH) for *POSTN* and *PECAM1* in human myocardial tissue confirming a distinct endocardial localization of *PECAM1 +/POSTN+* cells (Extended Data Fig. 5i).

Next, we modeled the spatially resolved expression of the two fibroblast subclusters fibroblast 1+2 (Fig. 2h) and identified several cytokines that explained the location of fibroblast 2. Interestingly, the model revealed that local expression of *IL13RA1*, linked to macrophage subtype 2 (Extended Data 5j), predicted the expression of the fibroblasts 2 marker *FBN1*, suggesting potential signaling between specific subpopulations of fibroblasts and macrophages in cardiac homeostasis (Fig. 2h, Extended Data Fig. 5j).

### Demarcation of the ischemic zone visualized by distinct gene expression and regulation

We next investigated the ischemic zone specimens of the acute human MI tissue that was collected 2-5 days after severe infarction (Fig. 3a-b, Extended Data Fig. 6a-d). snRNA-seq and snATAC-seq showed decreased cell capture rates due to the underlying biological process with ischemia-associated cell-death (Fig. 1h). We identified four distinct cell types in these datasets (Extended data Fig. 6e). From the spatial gene expression assays, we were able to generate data from this partially necrotic area with over 8,000 genes per spot in some locations (Extended Data Fig. 1e, IZ slides). Clustering of the spatial transcriptome data revealed a distinct pattern compared to the control heart with a demarcation of the infarcted tissue zone (Fig. 3a). We identified a core ischemic zone (spatial cluster 3) in the center of the specimen (Fig. 3a, right panel) surrounded by basophilic areas detectable in the hematoxylin and eosin-stained slide (Fig. 3a, left panel). This indicated the front of neutrophil granulocyte infiltration as the typical demarcation of nonperfused myocardial infarction. The front of neutrophil infiltration was detectable as cluster 8 surrounding the ischemic zone with expression of key neutrophil markers (*NCF1, ALOX5AP*) (Extended Data Fig. 6f). We further detected a strong gradient of *CXCL8* expression (cluster 2), a well-known neutrophil attractant chemokine (Fig. 3b). The severely damaged demarcated zone in the center of the specimen (cluster 3) was surrounded by concentric clusters of distinct gene expression (Fig. 3a-b). Of note, in this area the total number of genes detected was reduced compared to the other heart samples, in line with the expected celldeath in this ischemic area and explaining why the snRNA and snATAC data could only recover a few celltypes (Extended Data Fig. 1e). *TNNI3* (troponin) expression, a typical cardiomyocyte marker gene^9^, was identified in viable myocardium at the left edge of this specimen (Fig. 3b). Expression of the *NPPB* gene, coding for the widely used heart-failure biomarker brain natriuretic peptide (BNP)^18^, suggested a spatial zone of stressed surviving cardiomyocytes (Extended Data Fig. 6f). A zone of macrophage migration inhibitory factor (*MIF*) expression pointed towards strong macrophage infiltration and distinct zones of *COL3A1* expression indicated the presence of early activated fibroblasts (Fig. 3b, Extended Data Fig. 6f). We discovered a high importance of neighborhood expression of *S100A1* adjacent to the region of increased hypoxic signaling (Fig. 3c-d). Upregulation of *S100A1*, a calcium-binding protein that interacts with the contractile apparatus in cardiomyocytes^19^, has been shown to increase contractility in cardiomyocytes after myocardial infarction. The hypoxic signaling activity was highest in the area surrounding the necrotic zone, which could be explained by an enrichment of viable cells expressing hypoxia response genes (Fig. 3d). In addition to *CXCL8*, we also observed that the expression of *ANGPTL4*, a known facilitator of macrophage polarization in cardiac repair^20^ and *AREG*, a growth factor reported to be protective in cardiac ischemia^21^, were important predictors of distinct spatial zones (Fig. 3c). Furthermore, we found that the local expression of *S100A1* consistently predicts the expression of cardiomyocyte markers in both ischemic slides (Fig. 3c, Extended Data Fig. 7i).

**Fig. 3.**
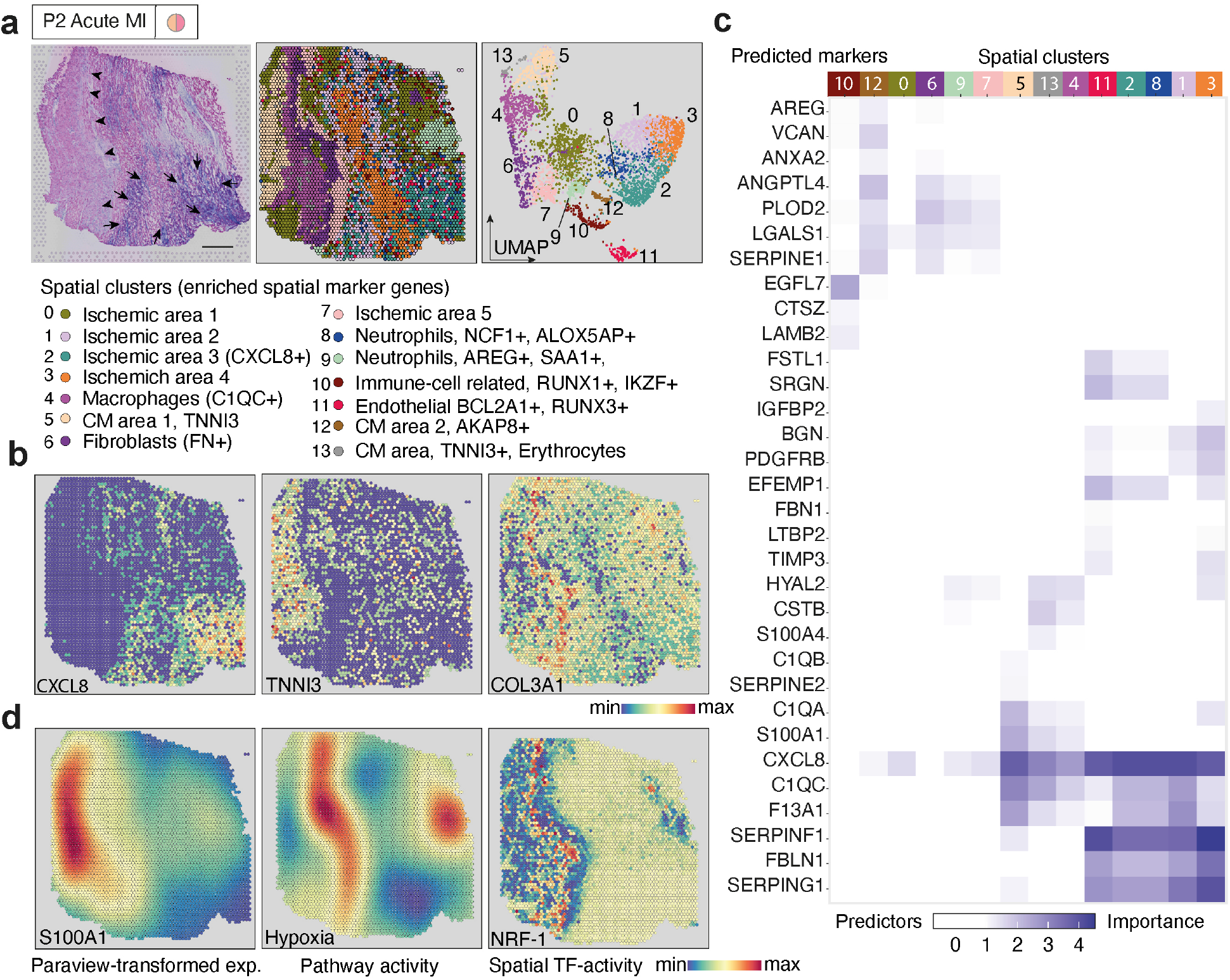
Spatial gene expression analysis of acute human myocardial ischemic tissue. **a.** Overview of the ischemic myocardial tissue (HE staining). Note the increments in blue staining (arrow) in the lower right area, indicating inflammatory cell infiltration. Clustering of the spatial detection spots resulted in 14 distinct clusters. Colors indicate individual clusters. UMAP embedding of spatial detection spots. Scale bar = 1mm. **b.** Scaled gene expression levels of CXCL8 (neutrophil recruitment), TNNI3 (cardiomyocytes), COL3A1 (fibroblasts). **c.** MISTy mean importances of the spatial expression of extracellular matrix and cytokine genes to explain the expression of marker genes of each cluster (for marker genes see Supplementary file 2). **d.** Paraview-transformed spatial expression (l = 10, see Methods) of S100A1 (left) and Hypoxia pathway activity (middle). Transcription factor (TF)-binding activity of NRF-1 mapped to each spot (right).

To explore the TFs associated with spatial regions, we mapped TF binding activity from snATAC-seq data to space using cell type scores. We observed locally increased NRF-1 binding activity in the area that also showed high hypoxia signaling (Fig. 3d) and collagen expression (Fig. 3b, Extended Data Fig. 6e). We next corroborated that the spatial TF binding activity corresponded to the expression of specific NRF-1 target genes (Extended Data Fig. 6h). NRF-1 regulates the expression of genes encoding respiratory chain subunits and other genes involved in mitochondrial function^22,23^, in line with cellular response to hypoxia.

The spatial transcriptomic data of the ischemic zone within the second acute MI specimen showed similar demarcation (Extended Data Fig. 7). Predictive features of these zones were conserved when compared to the remote area of the same heart specimen (Extended Data Fig. 8). Interestingly, we observed early signs of neo-angiogenesis in an ischemic area (cluster 2) with expression of *EGFL7* and *SOX4* and a spatial cardiomyocyte cluster that showed ER stress marker gene expression^24^ such as *HERPUD1* (cluster 4) (Extended Data Fig. 7a-d). In summary, the data indicated distinct spatial gene regulation in response to the ischemia associated cell-death with gene regulation driving the acute cardiac injury response.

### Spatially distinct cardiomyocyte subpopulations in the border zone

The border zone of the myocardial infarction is of particular interest, since spatial remodeling of this area is inextricably linked to the recovery of cardiac function^25^. Despite homogenous hematoxylin and eosin staining and UMI distribution across the slide (Fig. 4a and Extended Data Fig. 1e), we observed extensively heterogeneous spatial gene expression and identified a specific spatial transition zone in the center of the specimen (cluster 1) (Fig. 4a, Extended Data 9a-d). This zone separated distinct areas of gene expression in the upper left part from the lower right of the specimen (Fig. 4a). We identified 13 cell types from snRNA-seq (*n* = 6,081) and eight cell types from snATAC-seq (*n* = 3,101) (Fig. 4b-c, Extended Data Fig. 9a). Interestingly, both datasets identified two distinct cardiomyocyte populations (cardiomyocytes 1 and 2) (Fig. 4b-c). While cardiomyocytes 1 were solely located below the transition zone (injured area) cardiomyocytes 2 were primarily located above the transition zone (Fig. 4d). Differential gene expression analysis in the scRNA-seq data revealed significant upregulation of *NPPB, ANKRD1* and *MYO18B* in cardiomyocytes 1 (Fig. 4d, Extended data Fig. 9e). Both *NPPB* and *ANKRD1* have been reported to be upregulated in the border zone after MI.^26,27^ Pathway analysis of the spatial gene expression data indicated increased TGFβ activity within the injury area (lower right) while we identified homogeneous distribution of hypoxia activity (Fig. 4d, Extended Data Fig. 9f).

**Fig. 4.**
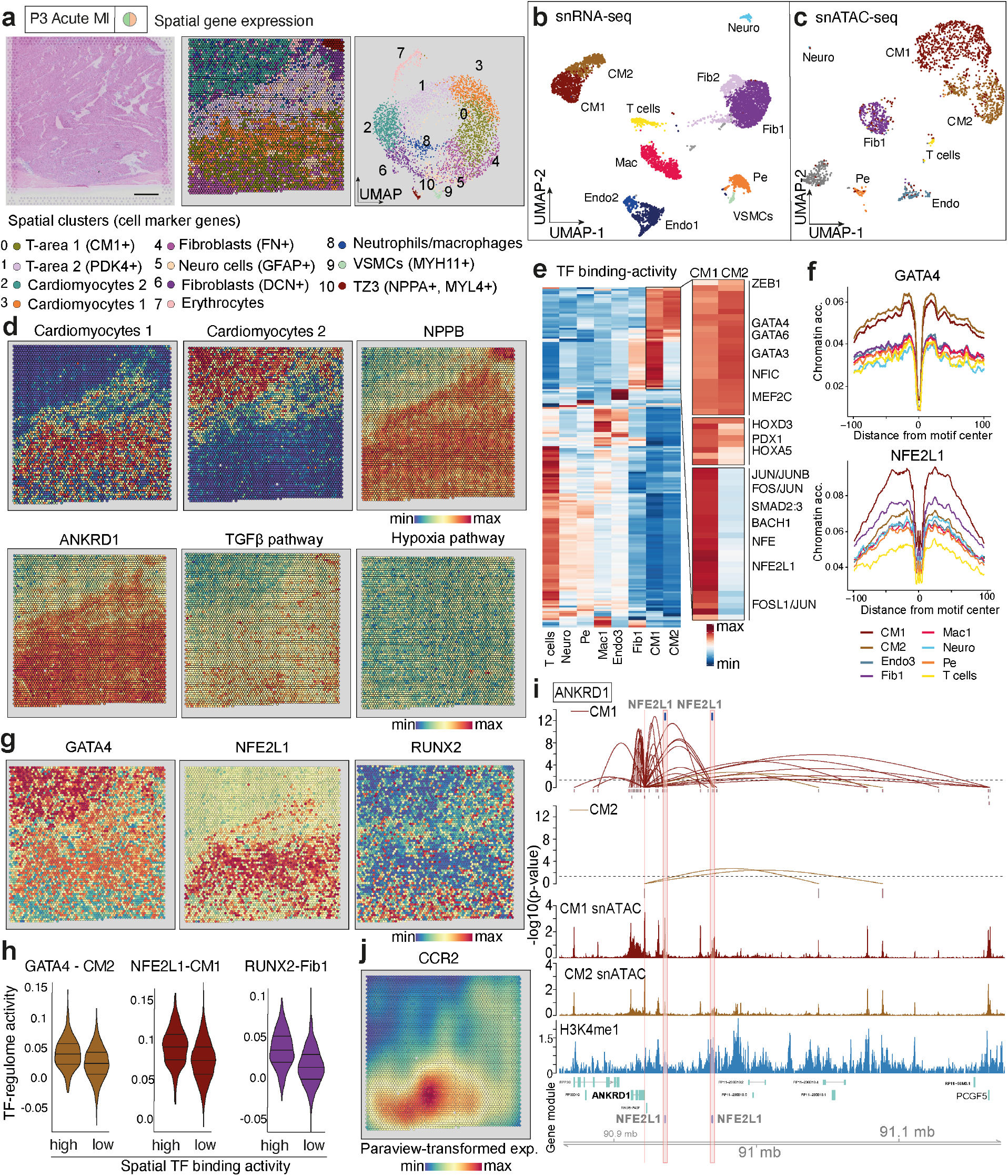
Gene regulatory changes of a cardiac borderzone delineated by multi-omic analysis. **a.** HE-staining of broderzone tissue showing homogenous tissue architecture. Unsupervised clustering of the spatial spots resulted in 10 clusters (left). UMAP embedding of the spatial spots (right). Scale bar = 1 mm. **b.** UMAP embedding of the snRNA-seq and **c.** snATAC-seq data. **d.** Cell-type score per spot by label transfer showing the distribution of cardiomyocyte 1 and cardiomyocyte 2. Scaled gene expression of the cardiac stress markers *NPPB* and *ANKRD1*. PROGENy’s^44,45^ Hypoxia and TGFβ pathway activity. Note that a gradient of hypoxia was not detected. **e.** Transcription factor (TF)-binding activity using HINT. Note the cardiomyocyte 1-specific activity of AP-1 complex TFs (JUN/FOS). **f.** Footprint profiles of GATA4 and NFE2L1 across different cell-types. Y-axis represents the average ATAC-seq signal around the TF-binding sites. **g**. Spatial distribution of GATA4, NFE2L1, and RUNX2 TF-binding activity. **h**. Violin plot comparing the summarized levels of expression using module scores of regulated genes of GATA4, NFE2L1, and RUNX2 of spots with high and low predicted TF-binding activities (Wilcoxon Rank Sum test, for GATA, NFE2L1 and RUNX2 p ≤ 9.1e-32). The 0.9 quantile separates high and low classes (high *n* = 467, low *n* = 4,194). **i.** Peak-to-gene links of ANKRD1 for cardiomyocyte 1 and cardiomyocyte 2. Each loop represents a putative link between ANKRD1 and a peak. Loop height represents the significance of the correlation and dash line represents threshold of significance (p = 0.05). ATAC-seq tracks were generated from pseudo-bulk chromatin profiles of cardiomyocyte 1 and cardiomyocyte 2. H3K4me1 ChIP-seq track of cardiomyocytes was obtained from an adult non-failing heart (Gilsbach, Ralf et al. 2018). Binding sites of NEF2L1 supported by ATAC-seq footprints are highlighted. **j.** Paraview-transformed spatial expression (l = 10) of CCR2.

Footprinting analysis of the snATAC-seq data revealed that both cardiomyocytes 1 and 2 had high binding activity of known cardiomyocyte lineage-specific TFs such as GATA4, MEF2C, and MYOD1, whereas cardiomyocytes 1 showed increased binding activity of NFE2L1 (Fig. 4e-f). NFE2L1 regulates a wide variety of cellular responses, several of which are related to important aspects of protection from ischemic stimuli^28^. Importantly, we observed increased binding activity of stress- and inflammation-associated TFs such as SMAD2/3, JUN, FOS (AP-1 signaling) in cardiomyocytes 1 compared to cardiomyocytes 2 (Fig. 4e). Interestingly, these gene regulatory programs of cardiomyocytes 1 were partly shared with cardiac fibroblasts (Fig. 4e). The overall switch of MEF2 to an AP-1 driven gene program has previously been described in the context of the border zone of murine MI^29^. Mapping of the TF binding activity into space indicates GATA4 TF binding activity in cardiomyocyte 2 location, whereas NFE2L1 and RUNX2 are associated with cardiomyocyte 1 and fibroblast 1 location, which we validated by analyzing the spatial expression of their cell-type specific target genes (Fig. 4g-h).

To further investigate *cis*-regulatory interactions differentiating these cardiomyocyte cell types, we detected peak-to-gene links by integrating snATAC- and snRNA-seq data (Extended data Fig. 10a)^30^. We next identified genes with the highest number of links specific to the cardiomyocyte 1 or 2 population (Extended data Fig. 10b). *ANKRD1* is among the top genes with significantly elevated peak-gene links in cardiomyocytes 1 compared to cardiomyocytes 2 (Extended data Fig. 10b). Interestingly, we detected two footprint-supported NFE2L1 binding sites in cardiomyocyte 1 specific peaks upstream of the *ANKRD1* promoter site (Fig. 4i). Additionally, high H3K4me1 signals in these regions supports enhancer function of these *cis*-regulatory elements in cardiomyocytes (Fig. 4i)^31^. These results indicate a direct regulation of *ANKRD1* by NFE2L1 in cardiomyocytes 1. Additionally, we identified footprints of NFE2L1 in enhancer regions of *MYO18B* and RUNX2 binding sites in enhancer regions of the *COL1A1* and *CREB3L2* gene (Extended data Fig.10c-e). Interestingly, we also observed that several cytokines were associated with the location of these cardiomyocyte subtypes (Extended Data Fig. 10f). After modeling the spatially resolved expression of cardiomyocyte cell marker genes, we observed that the local expression of *CCR2* predicted the expression of cardiomyocyte 1 marker genes in the injured zone of the specimen (Fig. 4j). CCR2^+^ resident cardiac macrophages have been reported as key regulators of an inflammatory response to myocardial injury^32^. Interestingly, *CCR2* was also of particular importance for predicting the localisation of fibroblast 1 together with TGFβ2 pathway activity suggesting potential crosstalk (Extended Data Fig. 10g). The second border-zone specimen overall showed similar findings (Extended Data Fig. 11).

In summary, the analysis indicated spatially distinct gene-expression and regulation in the borderzone with cardiomyocyte subsets that are specifically located within distinct regions of injury, inflammation and remodeling.

### Remodeling of the myocardium post-MI

Scar formation after MI is important for cardiac tissue integrity since failed scar formation can lead to ventricular rupture and death. One chronic remodeled specimen contained two visible scars, with one located in the center forming a U-shape (Fig. 5a, arrows) and a second one in the upper right corner of the specimen (Fig. 5a, arrowheads). Clustering of the spatial transcriptomics data revealed that the two scars aligned with distinct clusters (cluster 2 and cluster 9) and were separated by a larger area of endothelial cells reflecting neo-angiogenesis (Fig. 5a). We identified distinct subpopulations of fibroblasts and endothelial cells from snRNA-seq (*n* = 2,869) and snATAC-seq (*n* = 1,605) that showed an interesting spatial distribution in regard to the two scars (Fig. 5b-c, Extended Data Fig. 12a-c).

**Fig. 5.**
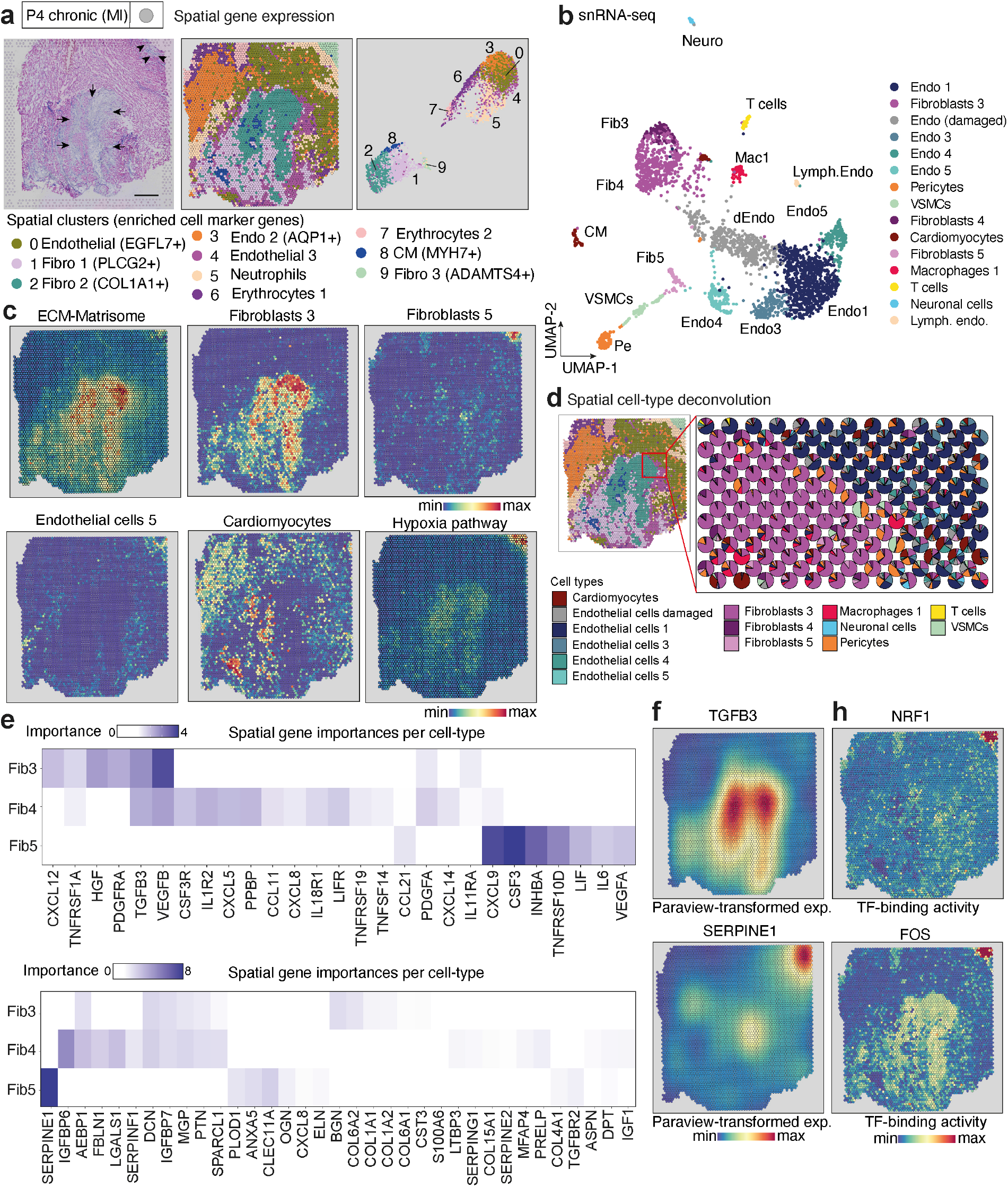
Temporal scar differences of fibrotic human heart. **a.** Overview of myocardial fibrotic tissue (HE-staining). Note the central U-shaped scar (arrows) and the top right corner scar (arrowheads). Clustering of the spatial spots resulted in 10 clusters. Cluster 2 and 9 aligned with observed scars. Scale bar = 1 mm. **b.** UMAP embedding of the snRNA-seq data with 15 cell types. Note that cardiomyocytes represent a minor fraction. **c.** Aggregated expression of extracellular matrix (ECM)-associated genes (top left). Cell-type score per spot by label transfer showing the distribution of cardiomyocytes, fibroblasts 5, fibroblasts 3, and endothelial cells. PROGENy’s^44,45^ hypoxia pathway activity (bottom right). Note the increased activity in the central U-shaped scar area and the enhanced activity in the right upper corner. **d.** Visualization of the distribution of cell-type scores at the interface of a fibrotic scar and adjacent endothelial cells. **e.** MISTy mean importances of the spatial expression of ECM, cytokine and cytokine receptor associated genes to explain the expression of marker genes of fibroblasts 5 and fibroblasts 3. For marker genes see Supplementary File 2. **f.** Paraview-transformed spatial expression (l = 10, see Methods) of TGFβ3 and SERPINE1. **h.** NRF1 and FOS transcription factor (TF)-binding activity mapped onto the spatial dataset from a.

Integration of snRNA and snATAC with spatial transcriptomics allowed us to increase the spatial resolution demonstrating the dominance of fibroblasts 3 within the central scar with a sharp border to a region that was dominated by endothelial cells 1 (Fig. 5d). This area was surrounded by an area with an increased abundance of pericytes also pointing towards neo-angiogenesis around the scar (Fig. 5d, Extended Data Fig. 12a). Interestingly, fibroblasts 5 were mainly present in the scar at the upper right corner of the slide surrounded by endothelial cells 5 (Fig. 5c). This scar showed less extracellular matrix (ECM) expression as compared to the central scar, which potentially represents a different stage of scar formation (Fig. 5d). Of note, the snRNA-seq and spatial transcriptomics data only identified a small population of cardiomyocytes (Fig. 5c-d) in line with the replacement scar formation after MI. Differential analysis of spatial features revealed increased JAK-STAT and TGFβ activity, both important pathways of fibrotic remodeling, in areas where fibroblasts 3 were enriched (spatial cluster 2) (Extended data Fig. 12e). The areas enriched with fibroblasts 5 also contained damaged endothelial cells and increased hypoxia pathway activity (Fig. 5d, Extended Data Fig. 12f). We observed that the local expression of *TGFβ3, PDGFRa* and *PDGFA* were highly important predictors of the presence of fibroblasts 3, which are abundant in the central scar area (Fig. 5e-f). *SERPINE1 (PAI-1*), which is associated with active scarring in tissues^33^, showed a high importance in cluster 9 and was associated with higher hypoxia and VEGF-A signaling (Fig. 5e-f, Extended data Fig. 12g). In contrast, the central scar area (spatial cluster 9) was also associated with higher binding activity of NRF-1 and the TF FOS (Fig. 5h). We further validated this using the parallel snATAC-seq data which suggest a higher binding activity of SMAD2/3 downstream in the TGFβ pathway (Extended data Fig. 12h). Altogether this data suggests a temporal difference between the two scar areas represented by the presence of a younger scar (upper right) with strong hypoxia signaling and a distinct fibroblast subpopulation and an older chronic large scar in the center with high matrix production involving FOS, SMAD2/3, NRF1 and RUNX1 (Extended Data Fig. 12h). This finding was also supported by higher spatial RUNX1 activity in fibroblasts 1 location of the second chronic dataset (Extended Data Fig. 13 a-j).

Since fibrotic response and scarring are the major feature of remodeling at late stage MI and are associated with cardiac stiffening, decreased systolic and diastolic function as well as triggering electric instability^34–36^, we next asked which mechanisms underlie fibroblast to myofibroblast differentiation in cardiac fibrosis.

### Trajectory analysis revealed RUNX1 as a regulator of myofibroblast differentiation in the human heart

To dissect mechanisms of myofibroblast differentiation we reclustered all fibroblasts from the integrated snRNA-seq dataset and identified 9 sub-clusters (Fig. 6a) suggesting an unappreciated heterogeneity of the cardiac stroma. Pseudotime analysis indicated a differentiation gradient of the integrated fibroblasts originating from cluster 3 and terminating in cluster 1 (Fig. 6a, right panel, Extended Data Fig. 14a). We have recently identified *SCARA5* as a marker for fibroblasts in the human kidney and demonstrated that *SCARA5^+^* fibroblasts are one origin of renal myofibroblasts.^37^ Therefore we used cells with the highest expression of *SCARA5* (cluster 3) as root cells in the pseudotime analysis. Periostin (*POSTN*) is a marker of myofibroblast differentiation that seems to be conserved between human and mouse and also plays a role in kidney fibrosis.^37^ Cluster 1 showed the highest expression of *POSTN, COL1A1* and *FN1* as well as collagen enrichment (NABA collagen score) compared to the *SCARA5^+^* cluster 3 (Fig. 6b-c, Extended Data Fig. 14a-b), which was indicative of a terminally differentiated myofibroblast population. The trajectory analysis inferred a differentiation trajectory from *SCARA5^+^* to *POSTN^+^* cells which was in line with our hypothesis. This data suggested downregulation of genes like *SCARA5* and *PCOLCE2* with increased expression of ECM genes and runt-related transcription factor 1 (RUNX1) during myofibroblast differentiation (Fig. 6d). To understand mechanisms of fibroblast to myofibroblast differentiation we next sorted genes, pathways and GO-terms (gene ontology) along this pseudotime trajectory which demonstrated late integrin signaling and ECM pathway enrichment consistent with fibroblast to myofibroblast differentiation (Extended Data Fig. 14b). Interestingly some of these findings, including reduced *SCARA5* and *PCOLCE2* expression in fibroblast to myofibroblast differentiation, appear to be conserved across different organs since we also observed this in the human kidney^37^. As suggested by our pseudotime analysis, we validated the presence of high *SCARA5^+^* expression in fibroblasts by co-staining with the pan-fibroblast/myofibroblast marker *COL15A1+* (Fig. 6 e, Extended Data Fig.14c-d) in human heart tissues. The direction of the trajectory was further confirmed by an accumulation of *POSTN+, COL1A1 +* myofibroblasts in fibrotic heart tissues of human heart failure patients (Fig. 6f, Extended Data Fig. 14e-f).

**Fig. 6.**
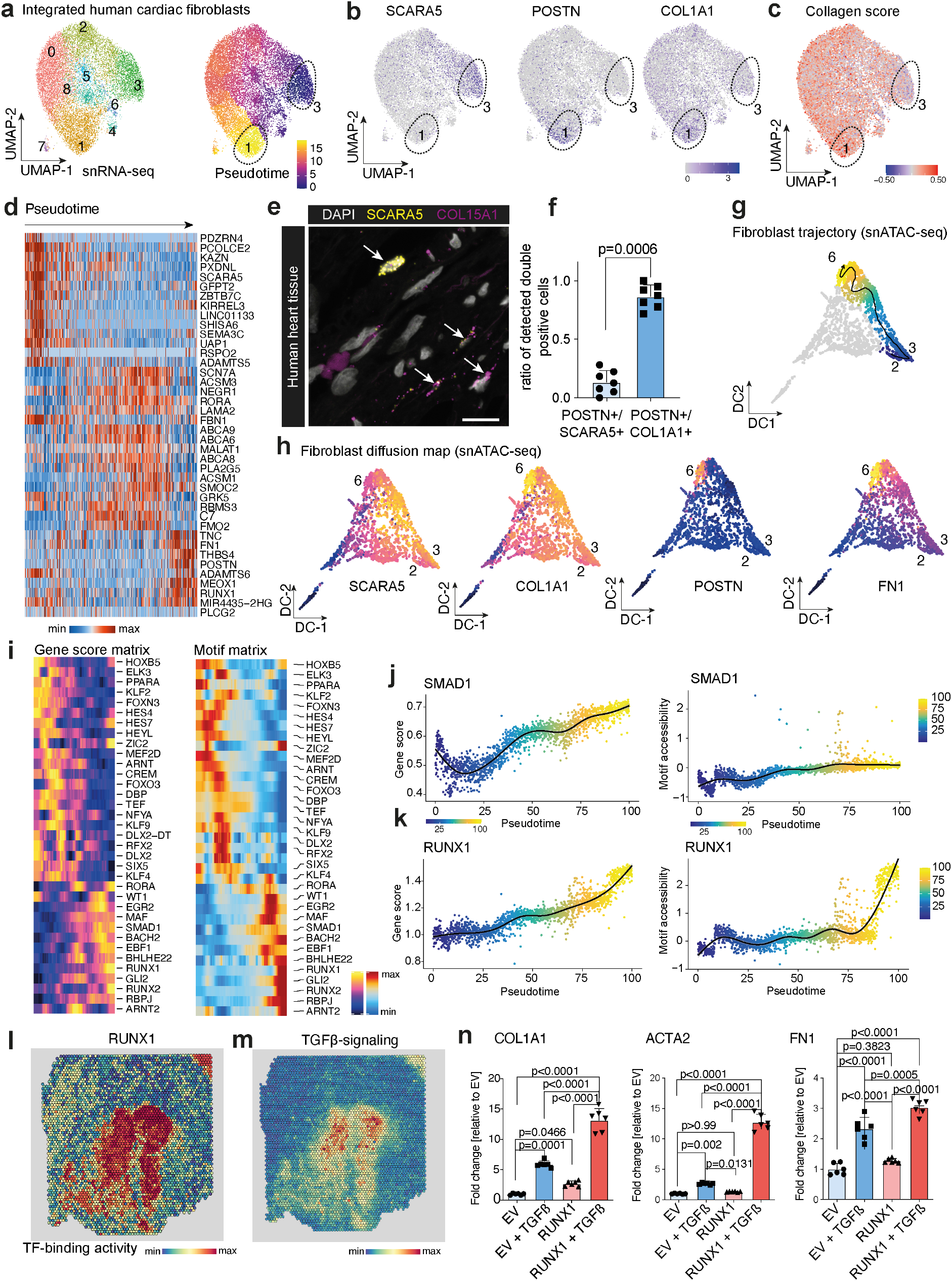
Trajectory analysis of cardiac myofibroblast differentiation. **a.** UMAP embedding of 14,637 fibroblasts from all snRNA-seq profiles (left), and Monocle3 pseudotime (right). **b.** Gene expression of *SCARA5, POSTN, COL1A1*, showing POSTN- and SCARA5-enrichment in clusters 1 and 3, respectively. **c.** Collagen score projected onto UMAP from a. **d**. Heatmap of differentially expressed genes across pseudotime from a. **e.** Representative in-situ hybridisation of *SCARA5* and *COL15A1* on human heart tissue. Scale bars: 20 μm. **f.** Quantification and comparison of SCARA5+/POSTN+ cells vs. POSTN+/COL1A1+ cells in human heart failure tissues (n=7). Mann-Whitney test. Error bars = S.D. **g.** Pseudotime trajectory of SCARA5^+^ and POSTN^+^ cells in snATAC-seq. The line represents a fitted trajectory across pseudotime. **h.** Scatter plot showing gene scores of SCARA5, POSTN, COL1A1 and FN1. **i.** Pseudotime heatmap showing gene scores (left) and TF motif activity (right) along the trajectory. **j.** Gene score (left) and motif accessibility (right) across the fibroblasts trajectory of SMAD1 and **k**. RUNX1. Each dot represents an individual pseudotime-ordered cell. **l.** RUNX1 motif activity in spatial data. **m.** TGFβ signaling in spatial data. **n.** Expression of *COL1A1, ACTA2* and *FN1* by RNA qPCR after *RUNX1* overexpression with and without TGFβ compared to empty vector (EV) (*n* = 6). One-way ANOVA followed by Bonferroni correction. Error bars = S.D.

We further verified this differentiation trajectory using the snATAC-seq data. Sub-clustering of all fibroblasts identified 11 distinct populations (Extended Data Fig. 14g). We next mapped the trajectory from snRNA-seq to snATAC-seq using ArchR^38^ (Fig. 6g-h). Importantly, we also observed cells with high accessibility of the *POSTN* and *SCARA5* promoter region in distinct ends of the diffusion map (Fig. 6g-h). To identify TFs that regulate this process, we integrated gene scores and TF motif activity along the trajectory and identified several TFs that showed a significant correlation between gene accessibility and motif activity along the trajectory (Fig. 6i, Extended Data Fig. 14h). We observed increased gene accessibility and motif activity for the Hedgehog transcription factor Gli2 along the fibroblast to myofibroblast differentiation trajectory (Extended Data Fig. 14i) which is in line with our reported role of Gli2 in regulating myofibroblast expansion^39^. We also observed an increase of SMAD1 activity and accessibility (Fig. 6j), which is a downstream TF of TGFβ signaling, one of the hallmark pathways of myofibroblast differentiation^40^. Interestingly, we observed that RUNX1 and RUNX2 also showed a gene accessibility and motif activity that increased from fibroblast to myofibroblast differentiation (Fig. 6k, Extended Data Fig. 14i). RUNX1 has been reported to physically interact with SMAD proteins and thus direct TGFβ signaling^40^. Our temporal TF analysis suggested SMAD1 was increasingly activated before RUNX1 (Fig. 6i) which is further supported by spatially co-localized RUNX1 TF binding activity and TGFβ signaling (Fig. 6l-m) and RUNX1/SMAD1 co-binding motif analysis (Extended Data Fig. 14j). The role of RUNX1 has been primarily studied in cardiomyocytes^41^ where deficiency protects against ischemic injury^42^. In the heart, the role of RUNX1 in non-myocytes is not defined in detail yet one recent study has reported the RUNX1 binding motif as a main motif in cardiac fibroblast-specific active enhancers^43^.

To validate the potential role of RUNX1 in myofibroblast differentiation we generated a human heart PDGFRβ^+^ fibroblast cell-line by FACS and lentiviral SV40LT and HTERT transduction (Extended Data Fig. 14k). Interestingly, lentiviral overexpression of RUNX1 in this cell-line increased TGFβ induced myofibroblast differentiation with increased expression of *COL1A1, ACTA2* and *FN1* (Fig. 6n). Therefore, our data indicates that RUNX1 is an important previously unappreciated driver of myofibroblast differentiation and acts by amplification of TGFβ signaling. Together with the known role of RUNX1 in cardiomyocytes this TF might be a very promising therapeutic target in cardiac remodeling after MI.

## Discussion

In multicellular organs, like the human heart, cellular function depends on the balance of interaction between neighbouring individual cell-types, which leads to tissue homeostasis. Single-cell technologies can profile the molecular heterogeneity of the different cell-types and their changes during disease. However, without spatial context it is unclear how these different cell types coordinate tissue functions. Here we provide a comprehensive resource map of the human heart at early and late stages after MI compared to control hearts (non-transplanted donor hearts), integrating spatial transcriptomics with single-cell gene expression and chromatin accessibility data. The use of spatial transcriptomics data allowed us to study links between cell lineages and functions that couldn’t be achieved from single-cell technologies alone.

We proposed a computational framework to integrate the multi-omics datasets of each patient. We related cell types across samples to achieve a high degree of consistency of the cell-type annotation. We increased the resolution of spatial transcriptomics by mapping cell-types to specific locations. Additionally, we provided a catalog of local biological processes that capture signaling and transcriptional regulatory events. Our findings provide detailed insights across cardiac cell-types and their response to ischemic injury of the human heart. We observed early demarcation of the infarcted area with regional influx of immunecell populations guided by regional expression of cytokines. The border zone surrounding the injured myocardium showed indeed a sharp border between injured and none-injured cell-types. Cardiomyocytes that differed by location with regard to the boundary revealed a distinct gene expression pattern and gene regulatory profile including enhancer accessibility. Late-stage remodeling after MI was driven by fibrosis with fibroblast to myofibroblast differentiation in distinct scars that were surrounded by areas of neoangiogenesis. Our data provides novel insights into myofibroblast differentiation in human hearts after MI, with distinct gene expression and gene regulatory programs driving this process including *RUNX1* as an amplifier of TGFβ signaling.

We envision that our work will serve as a reference for future studies integrating single cell (epi)genomics with spatial gene expression data. Furthermore, we believe that our data will facilitate the understanding of spatial gene expression and gene regulatory networks within the human myocardium and will be a resource for future studies that aim to understand the function of distinct cardiac cell types in cardiac homeostasis and ischemic disease.

## Acknowledgments

Funding: This work was supported by grants of the German Research Foundation (DFG: SFBTRR219 Project ID 322900939, CRU344,SCHN 1188/5-1 to RK and GE 2811/3-1 to IC), by a Grant of the European Research Council (ERC-StG 677448), a Grant of the State of North Rhine-Westphalia (Return to NRW), a Grant of the Else Kroener Fresenius Foundation (EKFS), the Dutch Kidney Foundation (DKF), TASKFORCE EP1805 all to RK, a grant of the Germany Society of Internal Medicine (DGIM) to CK, and a grant of the IZKF Faculty of Medicine at the RWTH Aachen University to IC. This work was also supported by the BMBF eMed Consortia Fibromap (to IC, RK and RKS). JSR and RORF are supported by Informatics for Life funded by the Klaus Tschira Foundation. Created with BioRender.com. 10xGenomics supported the project via the Visium challenge and provided free data generation of the Visium data.

## Author contributions

CK and RK planned and designed the study. CK, RR, ZL, MH, JT, HM, IC, FP, JSR, RK analyzed and interpreted the data. RR, ZL, MH, JT, IC, JSR designed the data analysis plan. CK, RR, ZL, MH, IC, JSR and RK wrote the manuscript and organized the figures. RSK and HM edited the manuscript and advised on data interpretation. HM organized patient tissue collection and biobanking and consented the patients. NK established the PDGFRb cell line from human hearts. SZ, XZ and CK carried out the overexpression studies. CK together with XZ and YX carried out all other single cell and imaging experiments. RMH and EMJB validated and sequenced the single cell libraries. IC, MC and ZL carried out primarily all snATAC-seq data analysis and its integration with snRNA-seq. RR, MH, JT and JSR carried out primarily the spatial gene expression and snRNA-Seq data analysis. CK, HM and RK initiated the study. All authors read and approved the manuscript.

## Supplementary Materials

- Supplementary File 1: Differential analysis of gene expression, transcription factor regulome-based activities (DoRothEA), pathway activities (PROGENy), and cell-type and ECM scores of each cluster of each spatial transcriptomics dataset.
- Supplementary File 2: Marker List for MISTy
- Supplementary File 3: Cell-type-specific TF binding activity
- Extended Data Table 1: Primer qPCR
- Extended Data Table 2: Clinical data of heart failure patients

## Data and Software availability

Custom scripts used in single cell data and spatial gene expression are available at: https://github.com/saezlab/visium_heart. Processed data from all human NGS-libraries that can be used as input to the scripts described above are available upon acceptance of the manuscript through the HCA.

## References

1. Sweeney, M., Corden, B. & Cook, S. A. Targeting cardiac fibrosis in heart failure with preserved ejection fraction: mirage or miracle? EMBO Mol. Med. e10865 (2020).

2. Kim, G.H., Uriel, N. & Burkhoff, D. Reverse remodelling and myocardial recovery in heart failure. Nat. Rev. Cardiol. 15, 83–96 (2018).

3. Wong, N.D. Epidemiological studies of CHD and the evolution of preventive cardiology. Nat. Rev. Cardiol. 11, 276–289 (2014).

4. Mendis, S., Puska, P., Norrving, B., Organization, W. H. & Others. Global atlas on cardiovascular disease prevention and control. (Geneva: World Health Organization, 2011).

5. Nabel, E. G. & Braunwald, E. A tale of coronary artery disease and myocardial infarction. N. Engl. J. Med. 366, 54–63 (2012).

6. Cung, T.-T. et al. Cyclosporine before PCI in Patients with Acute Myocardial Infarction. N. Engl. J. Med. 373, 1021–1031 (2015).

7. Niccoli, G. et al. Optimized Treatment of ST-Elevation Myocardial Infarction. Circ. Res. 125, 245–258 (2019).

8. Prabhu Sumanth D. & Frangogiannis Nikolaos G. The Biological Basis for Cardiac Repair After Myocardial Infarction. Circ. Res. 119, 91–112 (2016).

9. Litviňuková, M. et al. Cells of the adult human heart. Nature (2020) doi: 10.1038/s41586-020-2797-4.

10. Wang, L. et al. Single-cell reconstruction of the adult human heart during heart failure and recovery reveals the cellular landscape underlying cardiac function. Nat. Cell Biol. 22, 108–119 (2020).

11. Tucker, N. R. et al. Transcriptional and Cellular Diversity of the Human Heart. Circulation (2020) doi:10.1161/CIRCULATIONAHA.119.045401.

12. Li, Z. et al. Identification of transcription factor binding sites using ATAC-seq. Genome Biol. 20, 45 (2019).

13. Sun, S., Zhu, J. & Zhou, X. Statistical analysis of spatial expression patterns for spatially resolved transcriptomic studies. Nat. Methods 17, 193–200 (2020).

14. Kanisicak, O. et al. Genetic lineage tracing defines myofibroblast origin and function in the injured heart. Nat. Commun. 7, 12260 (2016).

15. Zhang, H. et al. Endocardium Minimally Contributes to Coronary Endothelium in the Embryonic Ventricular Free Walls. Circ. Res. 118, 1880–1893 (2016).

16. Tanevski, J., Gabor, A., Flores, R. O. R., Schapiro, D. & Saez-Rodriguez, J. Explainable multi-view framework for dissecting inter-cellular signaling from highly multiplexed spatial data. bioRxiv 2020.05.08.084145 (2020) doi:10.1101/2020.05.08.084145.

17. Tian, X. et al. Identification of a hybrid myocardial zone in the mammalian heart after birth. Nat. Commun. 8, 87 (2017).

18. Wei, C. M. et al. Natriuretic peptide system in human heart failure. Circulation 88, 1004–1009 (1993).

19. Most, P. et al. S100A1: a regulator of myocardial contractility. Proc. Natl. Acad. Sci. U. S. A. 98, 13889–13894 (2001).

20. Cho, D. I. et al. Antiinflammatory activity of ANGPTL4 facilitates macrophage polarization to induce cardiac repair. JCI Insight 4, (2019).

21. Koeppen, M. et al. Hypoxia-inducible factor 2-alpha-dependent induction of amphiregulin dampens myocardial ischemia-reperfusion injury. Nat. Commun. 9, 816 (2018).

22. Kelly, D. P. & Scarpulla, R. C. Transcriptional regulatory circuits controlling mitochondrial biogenesis and function. Genes Dev. 18, 357–368 (2004).

23. Virbasius, C. A., Virbasius, J. V. & Scarpulla, R. C. NRF-1, an activator involved in nuclear-mitochondrial interactions, utilizes a new DNA-binding domain conserved in a family of developmental regulators. Genes Dev. 7, 2431–2445 (1993).

24. Kokame, K., Agarwala, K. L., Kato, H. & Miyata, T. Herp, a new ubiquitin-like membrane protein induced by endoplasmic reticulum stress. J. Biol. Chem. 275, 32846–32853 (2000).

25. de Bakker, J. M. et al. Reentry as a cause of ventricular tachycardia in patients with chronic ischemic heart disease: electrophysiologic and anatomic correlation. Circulation 77, 589–606 (1988).

26. Hama, N. et al. Rapid Ventricular Induction of Brain Natriuretic Peptide Gene Expression in Experimental Acute Myocardial Infarction. Circulation vol. 92 1558–1564 (1995).

27. Mikhailov, A. T. & Torrado, M. The enigmatic role of the ankyrin repeat domain 1 gene in heart development and disease. Int. J. Dev. Biol. 52, 811–821 (2008).

28. Cui, M. et al. Dynamic Transcriptional Responses to Injury of Regenerative and Non-regenerative Cardiomyocytes Revealed by Single-Nucleus RNA Sequencing. Dev. Cell 53, 102–116.e8 (2020).

29. van Duijvenboden Karel et al. Conserved NPPB+ Border Zone Switches From MEF2-to AP-1– Driven Gene Program. Circulation 140, 864–879 (2019).

30. Corces, M. R. et al. The chromatin accessibility landscape of primary human cancers. Science 362, (2018).

31. Gilsbach, R. et al. Distinct epigenetic programs regulate cardiac myocyte development and disease in the human heart in vivo. Nat. Commun. 9, 391 (2018).

32. Bajpai Geetika et al. Tissue Resident CCR2- and CCR2+ Cardiac Macrophages Differentially Orchestrate Monocyte Recruitment and Fate Specification Following Myocardial Injury. Circ. Res. 124, 263–278 (2019).

33. Matsuo, S. et al. Multifunctionality of PAI-1 in fibrogenesis: Evidence from obstructive nephropathy in PAI-1–overexpressing mice. Kidney Int. 67, 2221–2238 (2005).

34. McDowell, K. S., Arevalo, H. J., Maleckar, M. M. & Trayanova, N. A. Susceptibility to arrhythmia in the infarcted heart depends on myofibroblast density. Biophys. J. 101, 1307–1315 (2011).

35. Miragoli, M., Salvarani, N. & Rohr, S. Myofibroblasts induce ectopic activity in cardiac tissue. Circ. Res. 101, 755–758 (2007).

36. Shinde, A. V. & Frangogiannis, N. G. Fibroblasts in myocardial infarction: a role in inflammation and repair. J. Mol. Cell. Cardiol. 70, 74–82 (2014).

37. Kuppe, C. et al. Decoding myofibroblast origins in human kidney fibrosis. Nature (2020) doi:10.1038/s41586-020-2941-1.

38. Granja, J. M. et al. ArchR: An integrative and scalable software package for single-cell chromatin accessibility analysis. Cold Spring Harbor Laboratory 2020.04.28.066498 (2020) doi: 10.1101/2020.04.28.066498.

39. Kramann, R. et al. Pharmacological GLI2 inhibition prevents myofibroblast cell-cycle progression and reduces kidney fibrosis. J. Clin. Invest. 125, 2935–2951 (2015).

40. Hanai, J.-I. et al. Interaction and Functional Cooperation of PEBP2/CBF with Smads. Journal of Biological Chemistry vol. 274 31577–31582 (1999).

41. Riddell, A. et al. RUNX1: an emerging therapeutic target for cardiovascular disease. Cardiovasc. Res. 116, 1410–1423 (2020).

42. McCarroll, C. S. et al. Runx1 Deficiency Protects Against Adverse Cardiac Remodeling After Myocardial Infarction. Circulation 137, 57–70 (2018).

43. Golan-Lagziel, T. et al. Analysis of rat cardiac myocytes and fibroblasts identifies combinatorial enhancer organization and transcription factor families. J. Mol. Cell. Cardiol. 116, 91–105 (2018).

44. Schubert, M. et al. Perturbation-response genes reveal signaling footprints in cancer gene expression. Nat. Commun. 9, 20 (2018).

45. Holland, C. H., Szalai, B. & Saez-Rodriguez, J. Transfer of regulatory knowledge from human to mouse for functional genomics analysis. Biochim. Biophys. Acta Gene Regul. Mech. 1863, 194431 (2020).

