## Extended Data Files for "Spatial multi-omic map of human myocardial infarction"

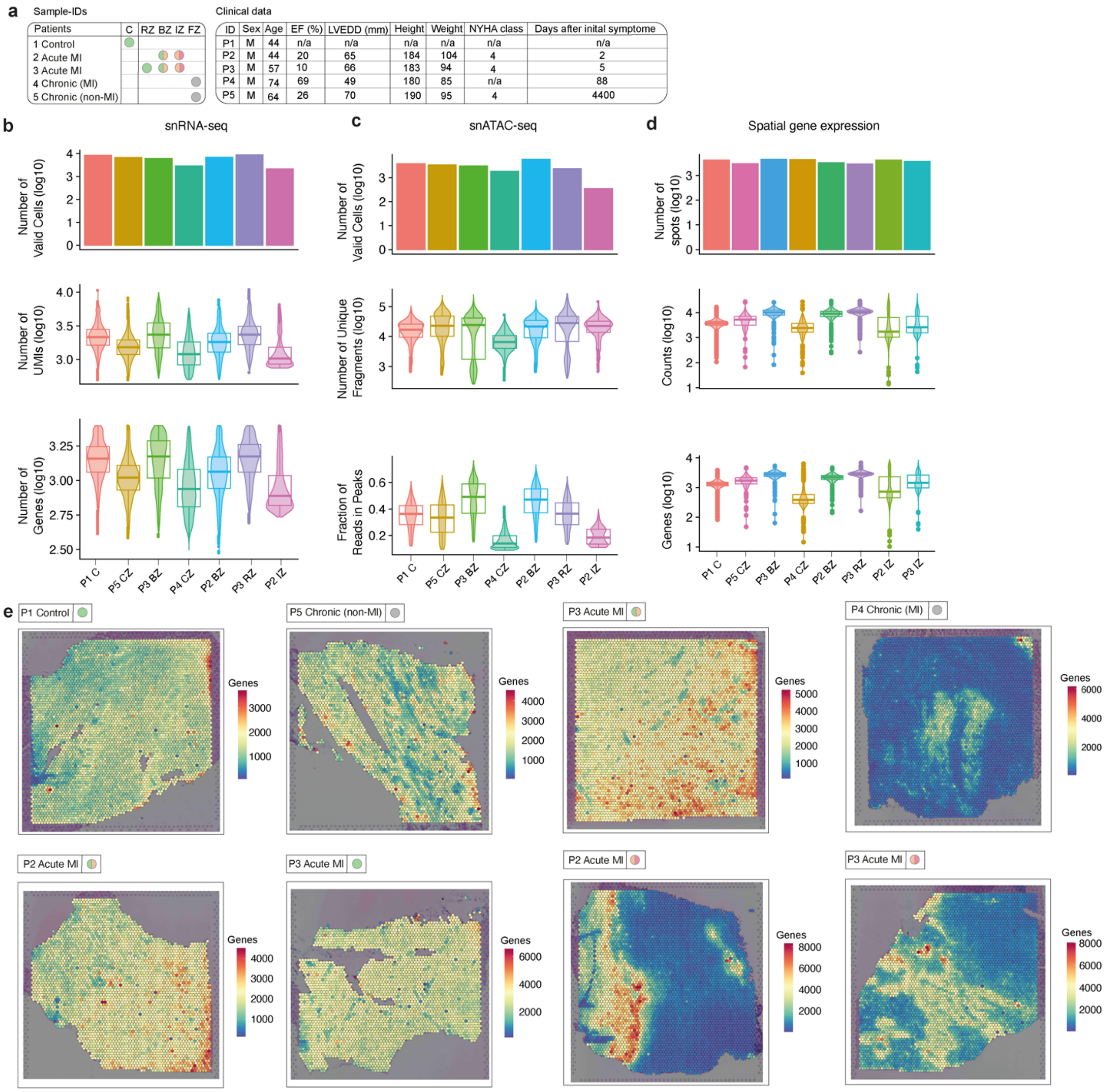


**Extended Data Fig. 1. Overview patient clinical data and dataset quality. a.** Sample ID and clinical data. C= control, RZ= remote zone, BZ= border zone, IZ= ischemic zone, FZ= fibrotic zone. EF refers to cardiac ejection fraction (in%), LVEDD to left ventricular end-diastolic diameter of the interventricular septum in mm, NYHA to New-York Heart Association classification of heart failure. **b.** Quality metrics of the snRNA-Seq datasets. Violin plots with horizontal lines indicate the median, the box indicates the span of the 25% to the 75% percentiles, whiskers extend to maximum 1.5x this interquartile range **c.** Quality metrics of the snATAC-Seq datasets. Violin plot same as in **b. d.** Quality metrics of the spatial gene expression datasets. Violin plot same as in **b. e.** Spatial distribution of number of detected genes per spot in the spatial gene expression datasets. Scale indicates number of genes per spot. Of note, the data quality of snRNA-seq from the ischemic sample (P3-IZ) was extremely low and we therefore only included the spatial transcriptomic data for this sample.

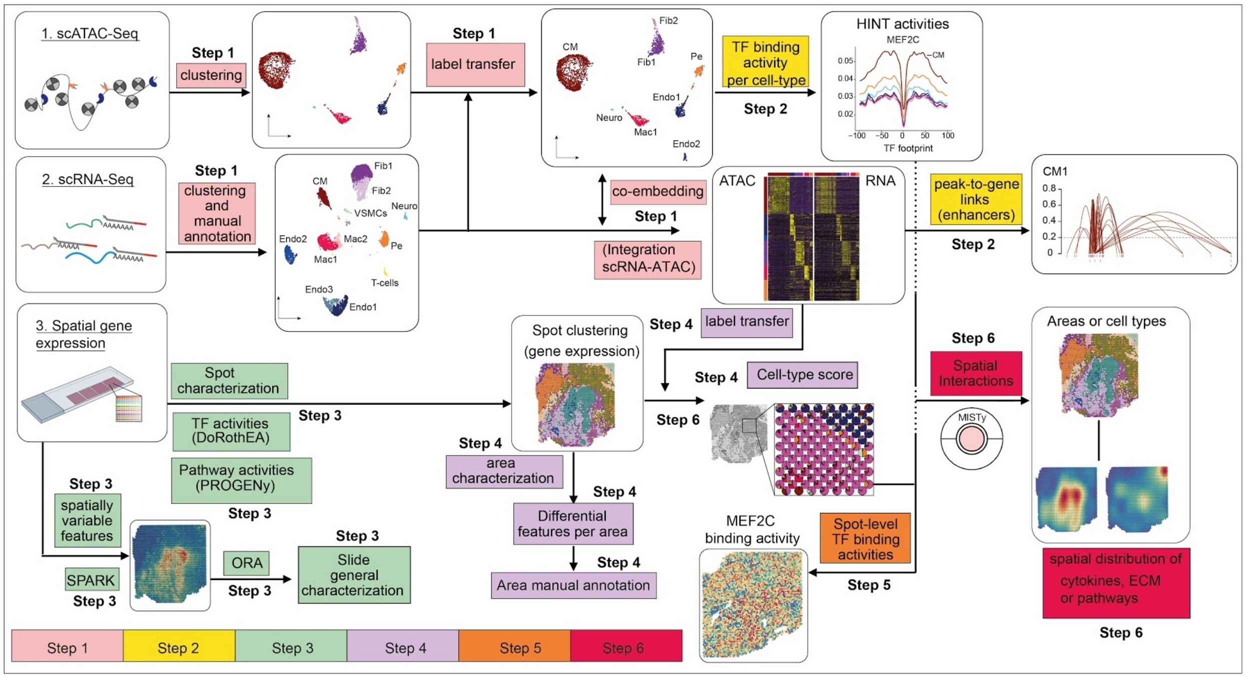


**Extended Data Fig. 2. Overview of computational workflow.** (***Step 1***) We clustered each dataset of snRNA- and snATAC-seq independently. We annotated the snRNA-seq data and used label transfer via Seurat for annotation of the snATAC-seq datasets. Single nuclear datasets were integrated using Seurat. (***Step 2***) We detected transcription factor (TF)-binding activities of specific cell-types with footprinting analysis on snATAC-seq data using HINT-ATAC. To uncover potential regulatory regions that control gene expression in different cell-types, we identified peak-to-gene links using the integrated snATAC- and snRNA-seq datasets. We used cell-specific footprints and peak-to-gene links to derive TF- and cell-specific regulomes. (***Step 3***) For the spatial data we used SPARK to identify spatially variable genes and overrepresentation analysis (ORA) for a general slide characterization. To functionally characterize each spot, we estimated the activities of TFs and signaling pathway activities using DoRothEA and PROGENy, respectively. (***Step 4***) Areas with similar patterns of expression were obtained using unsupervised clustering and differential expression analysis was used to manually annotate each area. We inferred the cell-type composition of each spot by transferring the cell-type annotations from their respective single nuclear datasets. (***Step 5***) TF-binding activity scores from snATAC datasets were mapped to the spatial location by considering the cellular composition of the spots. (***Step 6***) Finally, we studied spatial interactions using MISTy to estimate the importance of the local expression of putative extracellular matrix (ECM) proteins and cytokines, and pathway activities to the expression of marker genes of specific cell-types. For details see Methods.

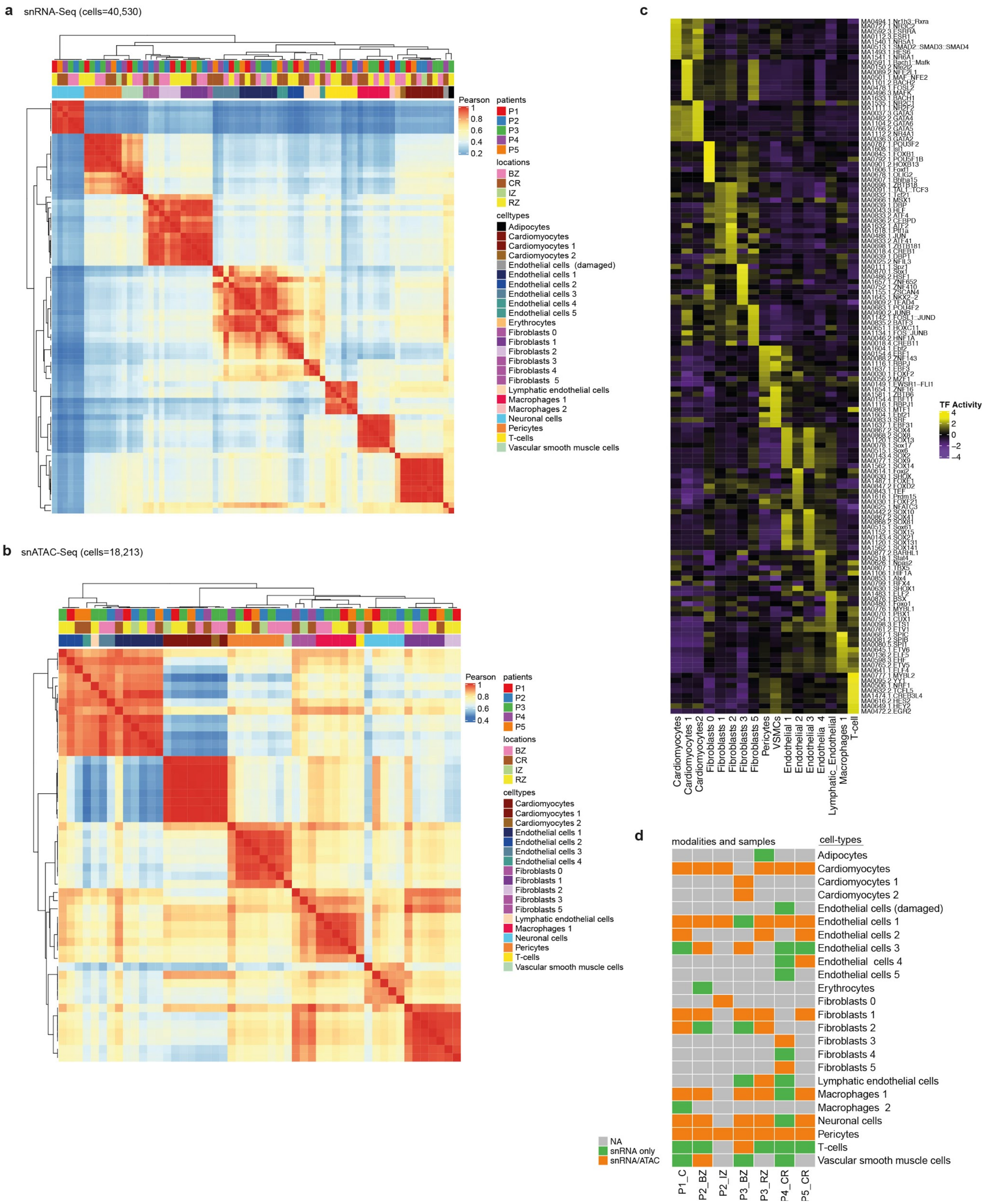
a
snRNA-Seq (cells=40,530)
Pearson
1
0.8
0.6
0.4
0.2
patients
P1
P2
P3
P4
P5
locations
BZ
CR
IZ
RZ
celltypes
Adipocytes
Cardiomyocytes
Cardiomyocytes 1
Cardiomyocytes 2
Endothelial cells (damaged)
Endothelial cells 1
Endothelial cells 2
Endothelial cells 3
Endothelial cells 4
Endothelial cells 5
Erythrocytes
Fibroblasts 0
Fibroblasts 1
Fibroblasts 2
Fibroblasts 3
Fibroblasts 4
Fibroblasts 5
Lymphatic endothelial cells
Macrophages 1
Macrophages 2
Neuronal cells
Pericytes
T-cells
Vascular smooth muscle cells
b
snATAC-Seq (cells=18,213)
Pearson
1
0.8
0.6
0.4
patients
P1
P2
P3
P4
P5
locations
BZ
CR
IZ
RZ
celltypes
Cardiomyocytes
Cardiomyocytes 1
Cardiomyocytes 2
Endothelial cells 1
Endothelial cells 2
Endothelial cells 3
Endothelial cells 4
Fibroblasts 0
Fibroblasts 1
Fibroblasts 2
Fibroblasts 3
Fibroblasts 5
Lymphatic endothelial cells
Macrophages 1
Neuronal cells
Pericytes
T-cells
Vascular smooth muscle cells
c
TF Activity
4
2
0
−2
−4
Cardiomyocytes
Cardiomyocytes 1
Cardiomyocytes2
Fibroblasts 0
Fibroblasts 1
Fibroblasts 2
Fibroblasts 3
Fibroblasts 5
Pericytes
VSMCs
Endothelial 1
Endothelial 2
Endothelial 3
Endothelia 4
Lymphatic_Endothelial
Macrophages 1
T-cell
d
modalities and samples
cell-types
Adipocytes
Cardiomyocytes
Cardiomyocytes 1
Cardiomyocytes 2
Endothelial cells (damaged)
Endothelial cells 1
Endothelial cells 2
Endothelial cells 3
Endothelial cells 4
Endothelial cells 5
Erythrocytes
Fibroblasts 0
Fibroblasts 1
Fibroblasts 2
Fibroblasts 3
Fibroblasts 4
Fibroblasts 5
Lymphatic endothelial cells
Macrophages 1
Macrophages 2
Neuronal cells
Pericytes
T-cells
Vascular smooth muscle cells
NA
snRNA only
snRNA/ATAC
P1_C
P2_BZ
P2_IZ
P3_BZ
P3_RZ
P4_CR
P5_CR

**Extended Data Fig. 3. Correlation heatmaps of integrated snRNA and snATAC datasets. a.** Heatmap with hierarchical clustering (average linkage) of all 7 integrated snRNA-seq datasets. P corresponds to the patient ID (see Extended Data Fig. 1a), BZ = borderzone, CR = control, IZ = ischemic zone, RZ = remote zone. **b.** Heatmap with hierarchical clustering of all 7 integrated snATAC-seq datasets. **c.** Heatmap showing transcription factor (TF)-binding activity per cell-type and sample. **d.** Heatmap showing cell-type distribution across patient samples and modalities. Colors refer to whether the cell-type is identified for a particular sample and modality.

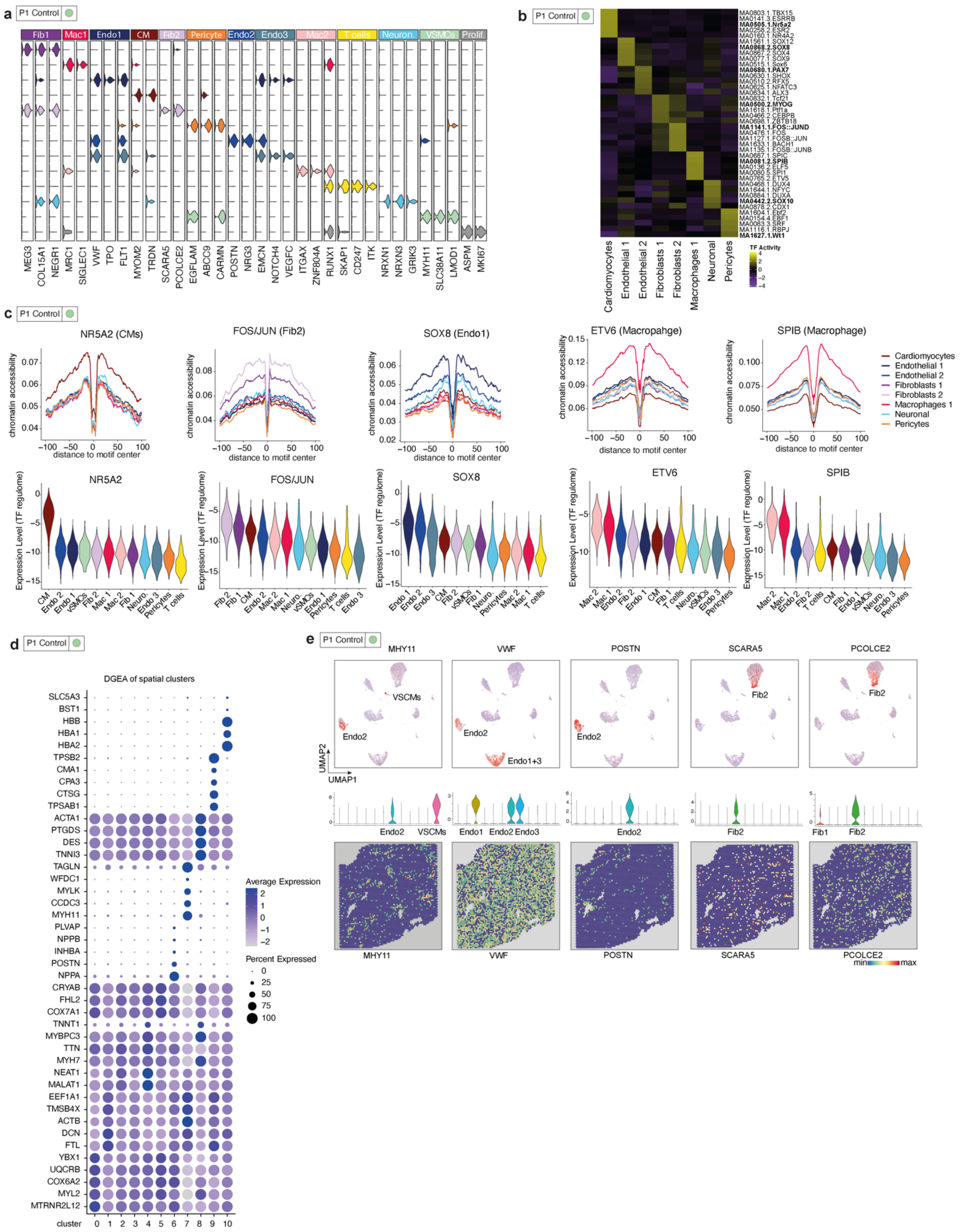
a
P1 Control
Fib1 Mac1 Endo1 CM Fib2 Pericyte Endo2 Endo3 Mac2 T cells Neuron. VSMCs Prolif.
MEG3 COL15A1 NEGR1 MRC1 SIGLEC1 VWF TPO FLT1 MYOM2 TRDN SCARA5 PCOLCE2 EGFLAM ABCC9 CARMN POSTN NRG3 EMCN NOTCH4 VEGFC ITGAX ZNF804A RUNX1 SKAP1 CD247 ITK NRXN1 NRXN3 GRIK3 MYH11 SLC38A11 LMOD1 ASPM MKI67
b
P1 Control
Cardiomyocytes Endothelial 1 Endothelial 2 Fibroblasts 1 Fibroblasts 2 Macrophages 1 Neuronal Pericytes
TF Activity
c
P1 Control
NR5A2 (CMs)
FOS/JUN (Fib2)
SOX8 (Endo1)
ETV6 (Macropahge)
SPIB (Macrophage)
chromatin accessibility
distance to motif center
Cardiomyocytes
Endothelial 1
Endothelial 2
Fibroblasts 1
Fibroblasts 2
Macrophages 1
Neuronal
Pericytes
NR5A2
FOS/JUN
SOX8
ETV6
SPIB
Expression Level (TF regulome)
d
P1 Control
DGEA of spatial clusters
SLC5A3 BST1 HBB HBA1 HBA2 TPSB2 CMA1 CPA3 CTSG TPSAB1 ACTA1 PTGDS DES TNNI3 TAGLN WFDC1 MYLK CCDC3 MYH11 PLVAP NPPB INHBA POSTN NPPA CRYAB FHL2 COX7A1 TNNT1 MYBPC3 TTN MYH7 NEAT1 MALAT1 EEF1A1 TMSB4X ACTB DCN FTL YBX1 UQCRB COX6A2 MYL2 MTRNR2L12
cluster 0 1 2 3 4 5 6 7 8 9 10
Average Expression
Percent Expressed
e
P1 Control
MHY11 VWF POSTN SCARA5 PCOLCE2
VSCMs Endo2 Endo1+3 Fib2
UMAP2 UMAP1
Endo2 VSCMs Endo1 Endo2 Endo3 Endo2 Fib2 Fib1 Fib2
MHY11 VWF POSTN SCARA5 PCOLCE2
min max

**Extended Data Fig. 4. Characterization of control heart sample. a.** Violin plots of cell-type marker gene expression from all cells of UMAP in Fig. 2A of all 13 cell subtypes. Cell types are color coded. **b.** Heatmap showing transcription factor (TF)- binding activity across all cell-types, as shown in Fig. 2. Note distinct TF binding activities between subtypes of fibroblasts and endothelial cells. **c.** Line plots showing ATAC-seq footprint profiles of NR5A2, TCF21, FOS/JUN, SOX8, ETV6, SPIB (upper panels). Y-axis represents the average ATAC-seq signal around the TF binding sites. Violin plots showing summarized levels of expression of TF-regulated genes per cell-type using module scores (lower panels). **d.** Heatmap showing the top 5 differentially expressed genes of each spatial cluster. **e.** UMAP embedding from the control heart sample. Shown are marker genes for distinct cell-types and corresponding violin plots and their spatial distributions.

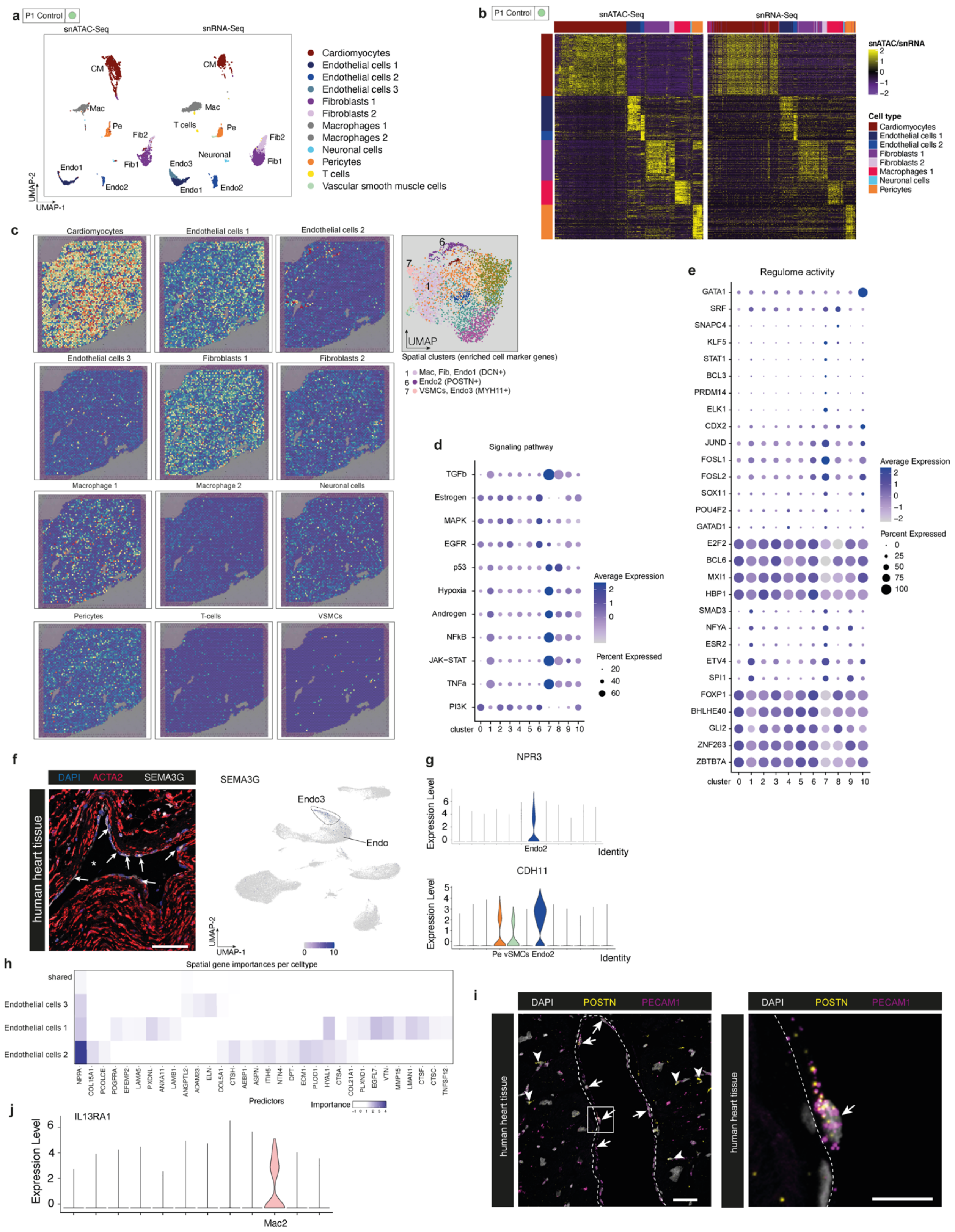

a
P1 Control
snATAC-Seq
snRNA-Seq
Cardiomyocytes
Endothelial cells 1
Endothelial cells 2
Endothelial cells 3
Fibroblasts 1
Fibroblasts 2
Macrophages 1
Macrophages 2
Neuronal cells
Pericytes
T cells
Vascular smooth muscle cells
b
snATAC/snRNA
Cell type
c
Spatial clusters (enriched cell marker genes)
1 Mac, Fib, Endo1 (DCN+)
6 Endo2 (POSTN+)
7 VSMCs, Endo3 (MYH11+)
d
Signaling pathway
Average Expression
Percent Expressed
e
Regulome activity
f
DAPI ACTA2 SEMA3G
human heart tissue
SEMA3G
g
NPR3
CDH11
Expression Level
Identity
h
Spatial gene importances per celltype
Predictors
Importance
i
DAPI POSTN PECAM1
human heart tissue
j
IL13RA1
Mac2

**Extended Data Fig. 5. Spatial characterization of the control heart sample. a.** Co-embedding of snATAC-seq (left) and snRNA-seq (right) cells in the same UMAP space. **b.** Heatmap showing peak-to-gene association by linking chromatin accessibility (left) and gene expression (right). Each column represents a cell from snATAC-seq (left) or snRNA-seq (right). Each row represents a peak (left) or gene (right). **c.** Cell-type score per spot by label transfer showing the distribution of distinct cell-types. Coverage and mean expression (activity or score) per cluster in spatial transcriptomics **d.** PROGENy's signaling pathway activities. **e.** The top 5 differentially active transcription factors (TFs) using DoRothEA. **f.** Immunofluorescent imaging of SEMA3G on human myocardial tissue, a marker of endothelial cell 3 and ACTA2. **g.** Violin plots showing the expression of *NPR3* and *CDH11* across all cell-types. **h.** MISTy mean importances of the spatial expression of extracellular matrix (ECM)- and cytokine-associated genes to explain the expression of marker genes of Endothelial cells 1-3. **i.** In-situ hybridisation of POSTN and PECAM1 on human myocardial tissue. Scale bars: 10 μm. **j.** Violin plot showing expression of *IL13RA1* in Mac2.

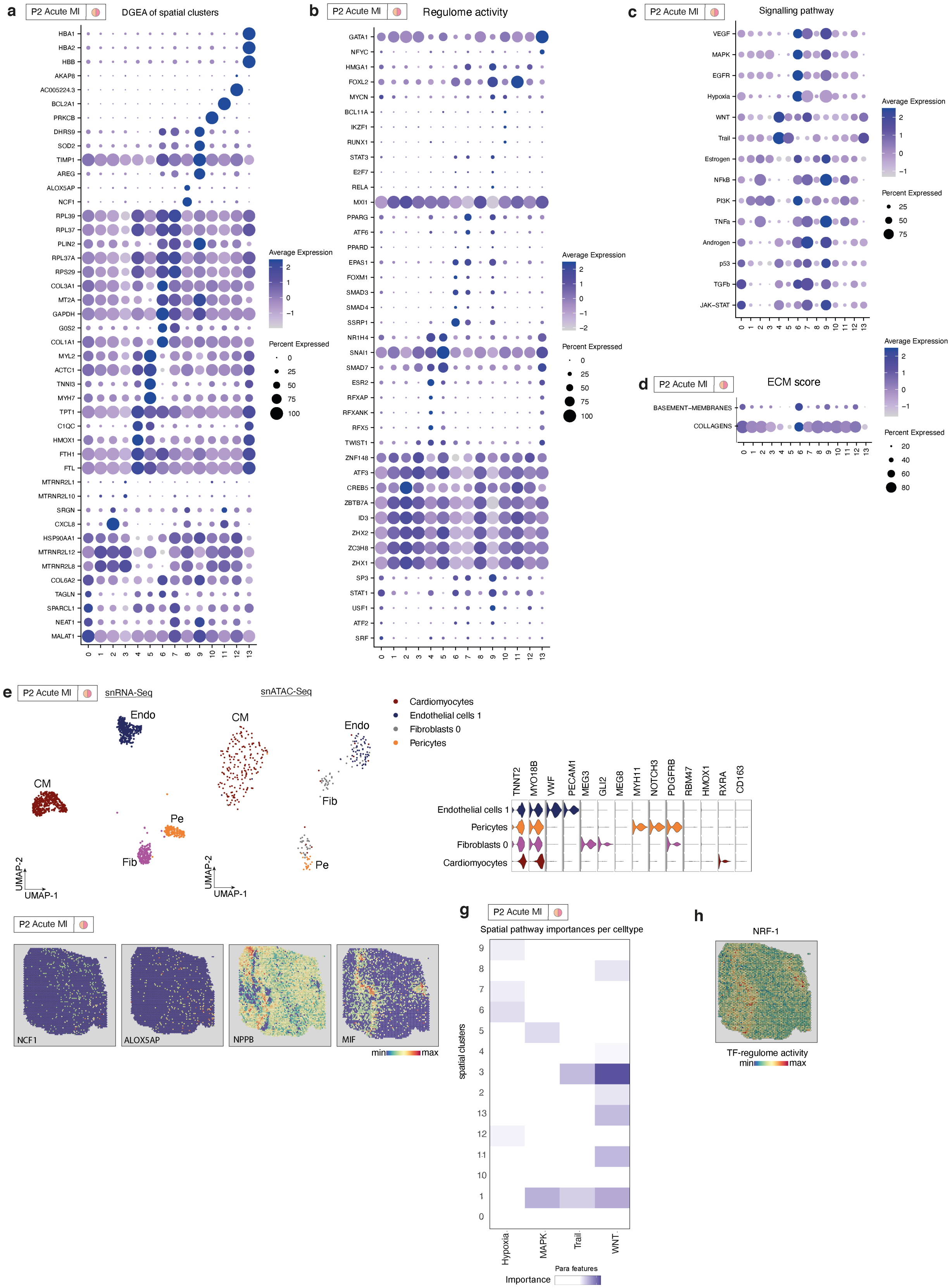

a
P2 Acute MI
DGEA of spatial clusters
b
Regulome activity
c
Signalling pathway
d
ECM score
e
snRNA-Seq
snATAC-Seq
Cardiomyocytes
Endothelial cells 1
Fibroblasts 0
Pericytes
f
NCF1
ALOX5AP
NPPB
MIF
g
Spatial pathway importances per celltype
h
NRF-1
TF-regulome activity

**Extended Data Fig. 6 Characterization of acute myocardial infarction sample.**

**a.** The top 5 differentially expressed genes of each cluster. **b.** The top 5 differentially active TFs using DoRothEA. **c.** PROGENy's signaling pathway activities and **d.** ECM scores (See Methods). **e.** UMAP embedding of single nuclei transcriptomes and open chromatin profiles from the ischemic myocardial tissue. **f.** Spatial distribution of NCF1, ALOX5AP, NPPB and MIF across the detection area**. g.** MISTy's mean importances of PROGENy's pathway activities (paraview) to explain the expression of markers across clusters. **h.** Summarized levels of expression of NRF-1 regulated genes using module scores (see Methods).

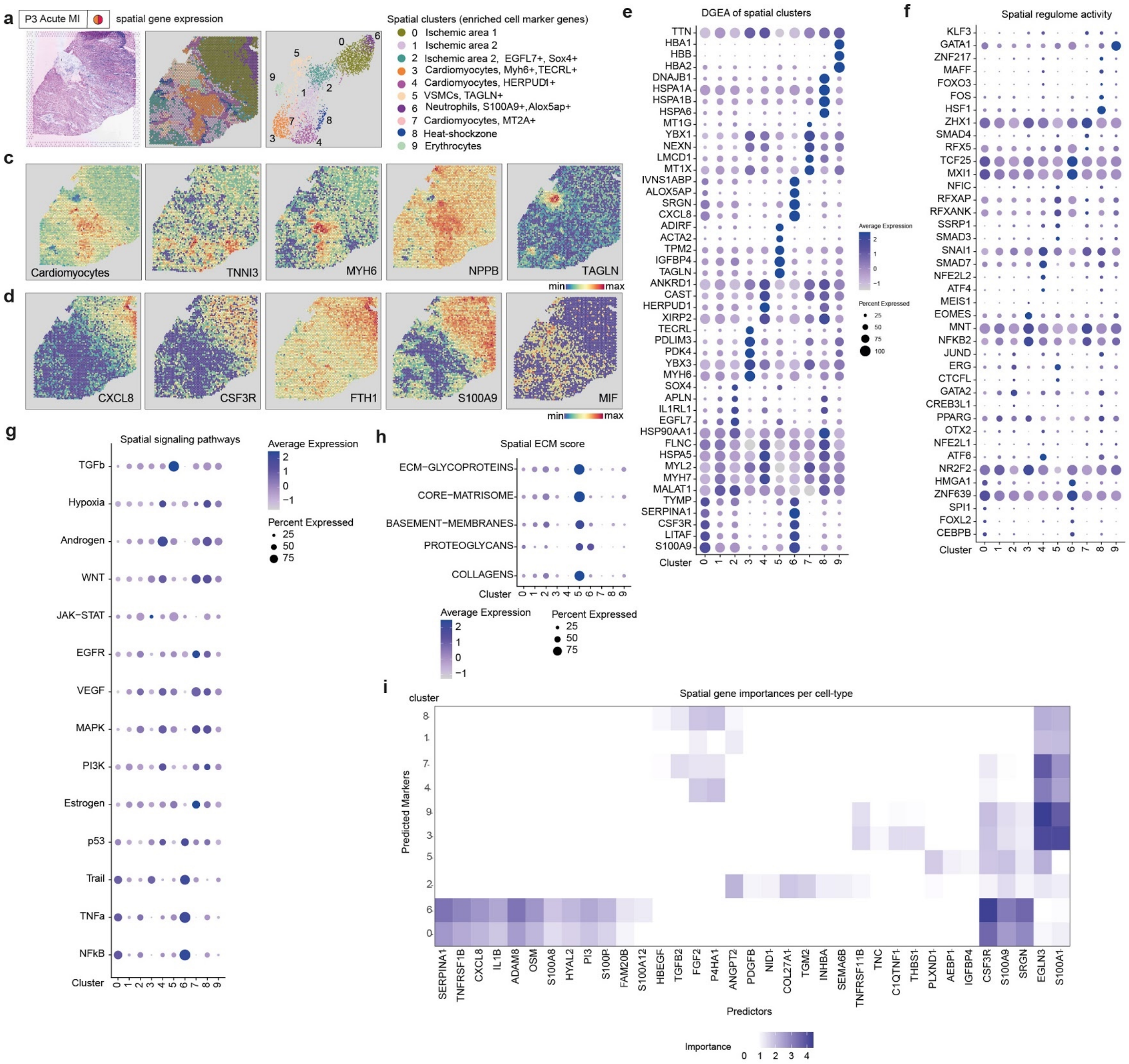

**Extended Data Fig. 7 Spatial muti-omic characterization of acute myocardial infarction sample. a.** Overview of the second ischemic myocardial tissue (HE staining). Note basophilic staining in the upper right corner of the detection slide. **b.** Clustering of the spatial detection spots resulted in 10 clusters. UMAP embedding of the spatial detection spots. **c.** Spatial distribution of cardiomyocytes (based on cell type scores) and the expression of TNNI3, MYH6, NPPB, and TAGLN. **d.** Spatial distribution of the expression of CXCL8, CSF3R, FTH1, S100A9, and MIF. Coverage and mean expression (activity or score) per cluster in spatial transcriptomics of: **e**. The top 5 differentially expressed genes of each cluster, **f.** The top 5 differentially active TFs using DoRothEA, **g.** PROGENy's signaling pathway activities, and **h.** ECM scores (See Methods). **i.** MISTy's mean importances of the spatial expression (paraview) of ECM genes and cytokines to predict the expression of markers across clusters.

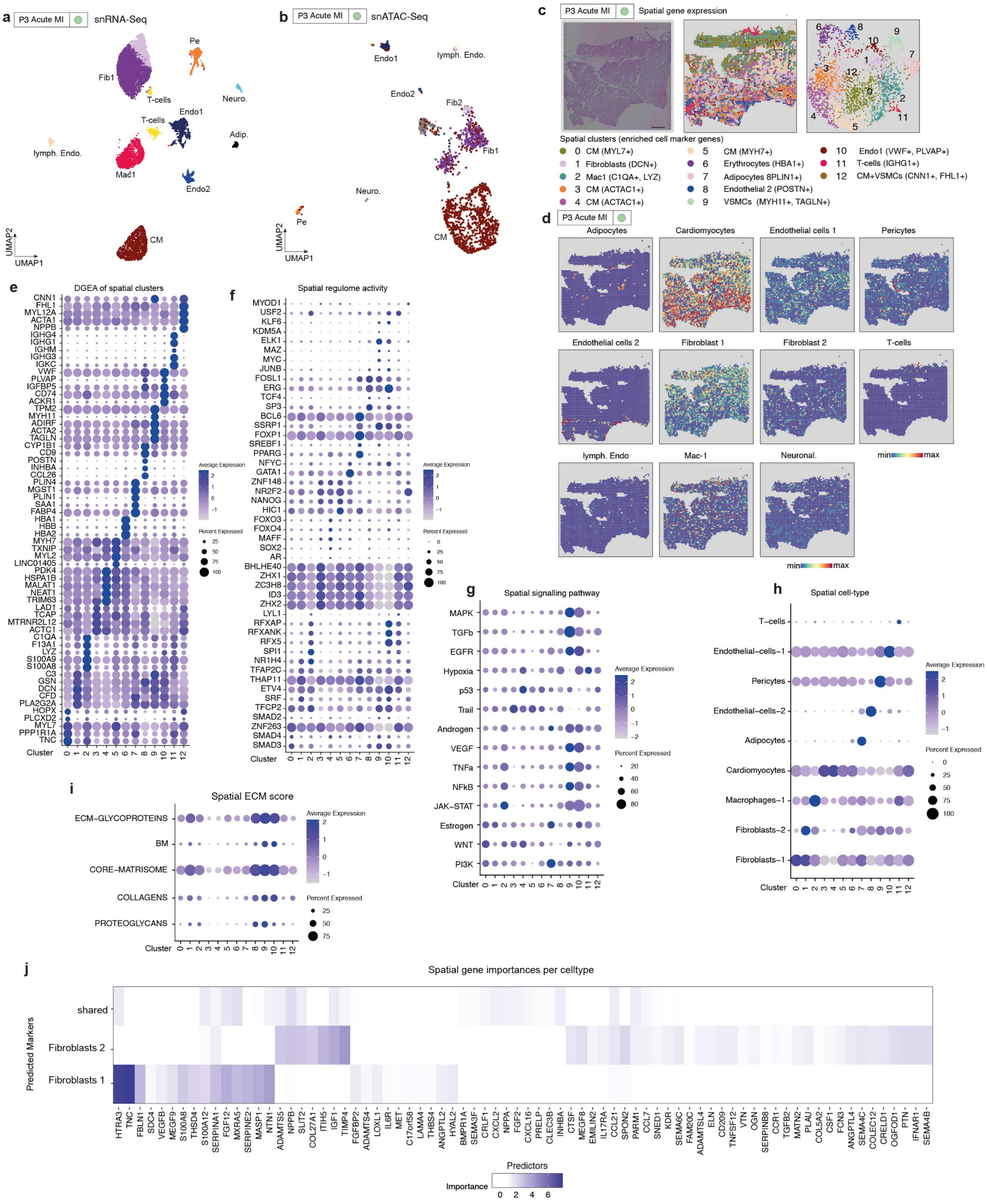

a
P3 Acute MI
snRNA-Seq
Pe
Fib1
T-cells
Neuro.
Endo1
T-cells
Adip.
lymph. Endo.
Mac1
Endo2
CM
UMAP2
UMAP1
b
P3 Acute MI
snATAC-Seq
Endo1
lymph. Endo.
Endo2
Fib2
Fib1
Neuro.
Pe
CM
UMAP2
UMAP1
c
P3 Acute MI
Spatial gene expression
Spatial clusters (enriched cell marker genes)
0 CM (MYL7+)
1 Fibroblasts (DCN+)
2 Mac1 (C1QA+, LYZ)
3 CM (ACTAC1+)
4 CM (ACTAC1+)
5 CM (MYH7+)
6 Erythrocytes (HBA1+)
7 Adipocytes 8PLIN1+)
8 Endothelial 2 (POSTN+)
9 VSMCs (MYH11+, TAGLN+)
10 Endo1 (VWF+, PLVAP+)
11 T-cells (IGHG1+)
12 CM+VSMCs (CNN1+, FHL1+)
d
P3 Acute MI
Adipocytes
Cardiomyocytes
Endothelial cells 1
Pericytes
Endothelial cells 2
Fibroblast 1
Fibroblast 2
T-cells
lymph. Endo
Mac-1
Neuronal.
min
max
e
DGEA of spatial clusters
Average Expression
Percent Expressed
Cluster
f
Spatial regulome activity
Average Expression
Percent Expressed
Cluster
g
Spatial signalling pathway
MAPK
TGFb
EGFR
Hypoxia
p53
Trail
Androgen
VEGF
TNFa
NFkB
JAK–STAT
Estrogen
WNT
PI3K
Cluster
h
Spatial cell-type
T–cells
Endothelial–cells–1
Pericytes
Endothelial–cells–2
Adipocytes
Cardiomyocytes
Macrophages–1
Fibroblasts–2
Fibroblasts–1
Cluster
i
Spatial ECM score
ECM–GLYCOPROTEINS
BM
CORE–MATRISOME
COLLAGENS
PROTEOGLYCANS
Cluster
j
Spatial gene importances per celltype
Predicted Markers
shared
Fibroblasts 2
Fibroblasts 1
Predictors
Importance

**Extended Data Fig. 8 Characterization of acute myocardial infarction sample remote zone. a.** UMAP embedding of the snRNA-Seq and **b.** snATAC-seq data. **c.** Clustering of the spatial spots resulted in 13 clusters (left and middle panel). UMAP embedding of the slide spots (right panel). **d.** Cell-Type score per spot by label transfer. Coverage and mean expression (activity or score) per cluster in spatial transcriptomics of: **e**. The top 5 differentially expressed genes of each cluster, **f.** The top 5 differentially active TFs using DoRothEA, **g.** PROGENy's signaling pathway activities, **h**. Cell-type scores as calculated with label transfer and **i.** ECM scores (See Methods). **j.** MISTy's mean importances of the spatial expression (paraview) of ECM genes and cytokines to predict the expression of marker genes across different fibroblast subtypes.

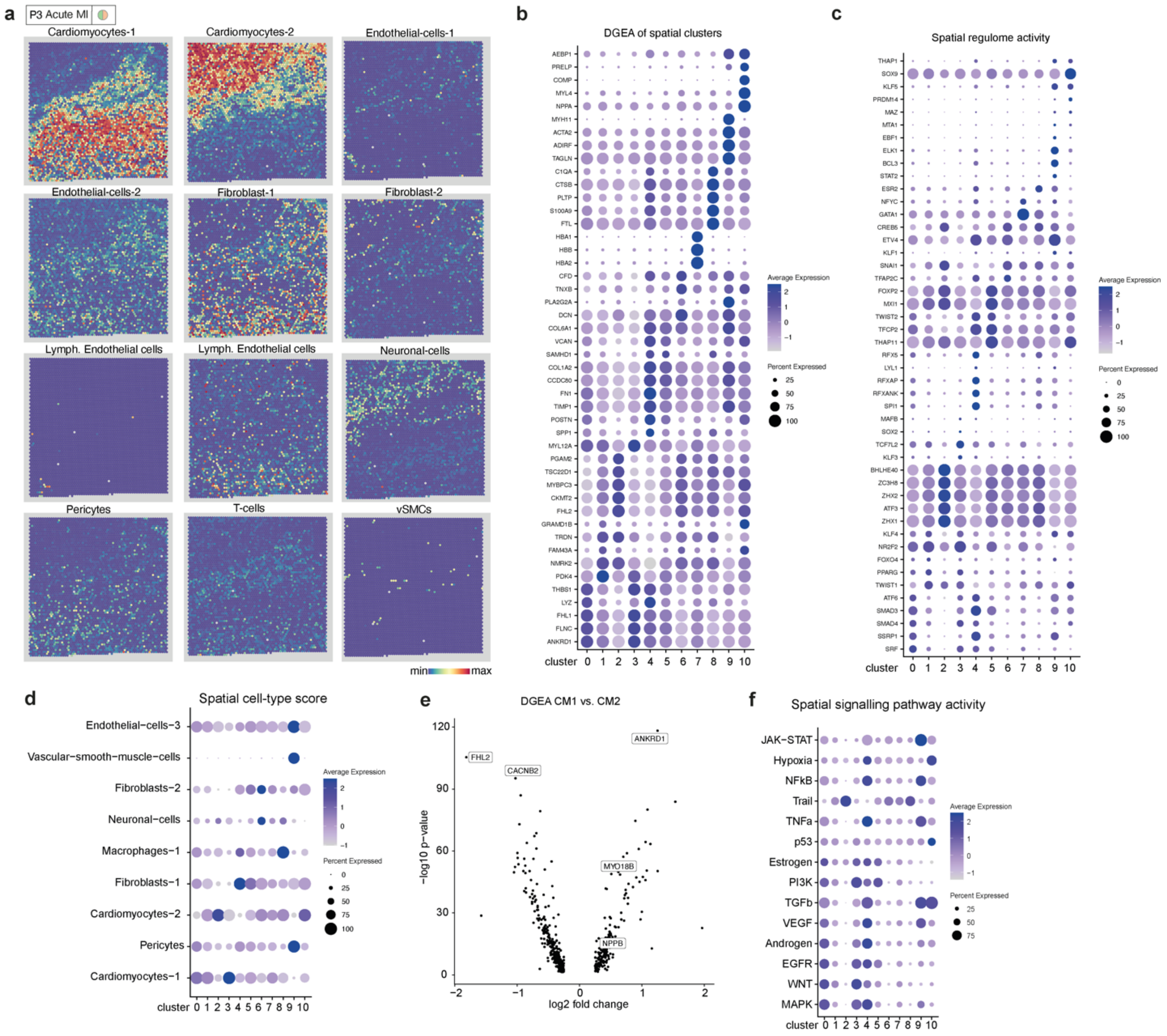

**Extended Data Fig. 9 Characterization of the borderzone of an acute myocardial infarction. a.** Cell-type score per spot by label transfer. Coverage and mean expression (activity or score) per cluster in spatial transcriptomics of: **b.** The top 5 differentially expressed genes of each cluster, **c.** The top 5 differentially active TFs using DoRothEA. **d.** Cell-type scores as calculated with label transfer (See Methods). **e.** Volcano plot showing differentially expressed genes between cardiomyocytes 1 (CM1) and cardiomyocytes 2 (CM2) from the snRNA-seq data. **f.** PROGENy's signaling pathway activities of spatial clusters.

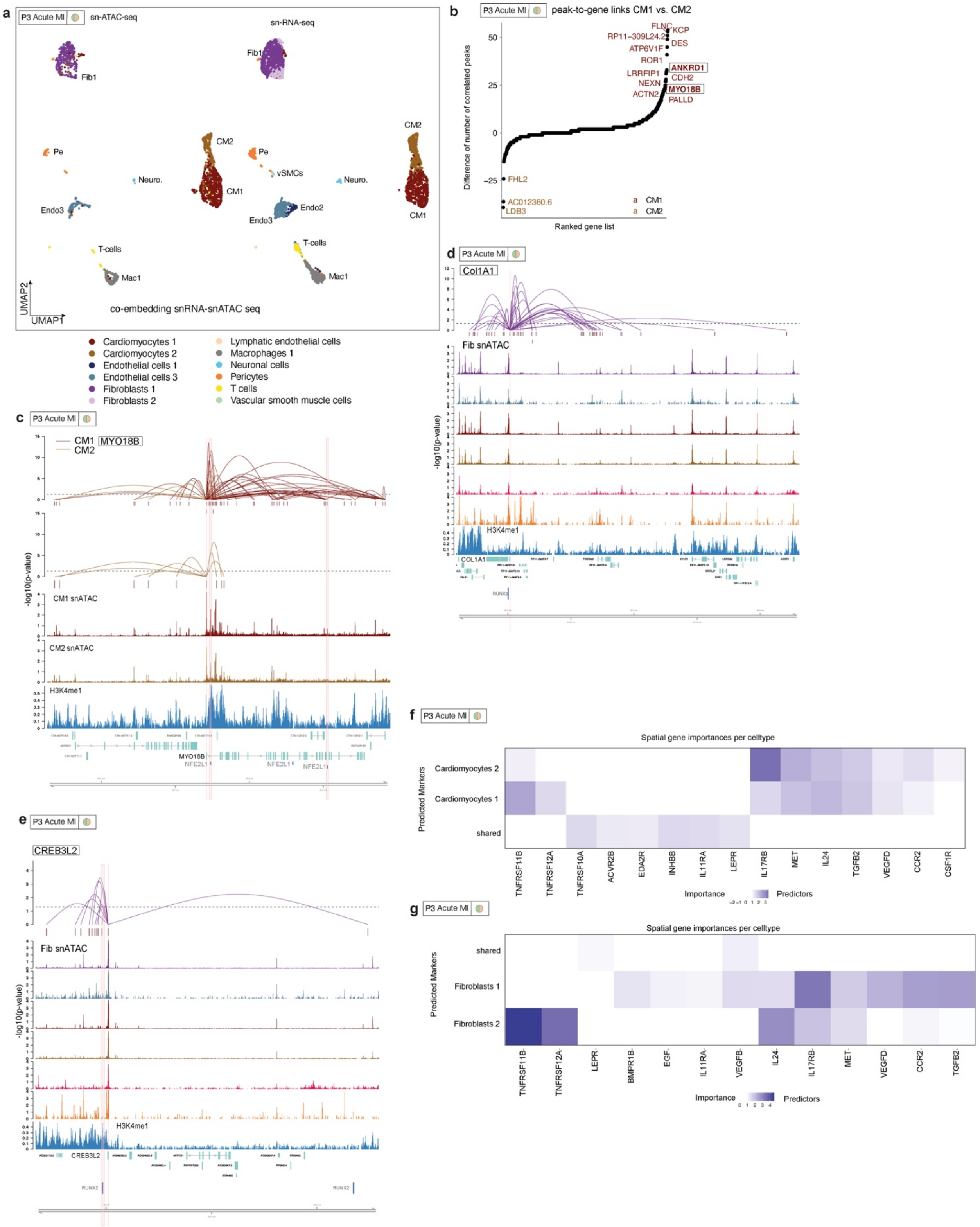
a
P3 Acute MI
sn-ATAC-seq
sn-RNA-seq
co-embedding snRNA-snATAC seq
Cardiomyocytes 1
Cardiomyocytes 2
Endothelial cells 1
Endothelial cells 3
Fibroblasts 1
Fibroblasts 2
Lymphatic endothelial cells
Macrophages 1
Neuronal cells
Pericytes
T cells
Vascular smooth muscle cells
b
peak-to-gene links CM1 vs. CM2
Difference of number of correlated peaks
Ranked gene list
c
MYO18B
CM1 snATAC
CM2 snATAC
H3K4me1
d
Col1A1
Fib snATAC
e
CREB3L2
f
Spatial gene importances per celltype
Predicted Markers
Importance
Predictors
g
Spatial gene importances per celltype

**Extended Data Fig. 10 Characterization of cis-regulatory elements and spatial gene importances in the myocardial borderzone. a.** UMAP demonstrates the co-embedding of snATAC-seq (left) and snRNA-seq (right) data into common low dimensional space. **b.** Ranked differentially expressed gene list based on the differences of number of associated peaks between cardiomyocytes 1 and cardiomyocytes 2. **c.** Peak-to-gene links of MYO18B for cardiomyocytes 1 and cardiomyocytes 2. Each loop represents a putative link between ANKRD1 and a peak. Loop height represents the significance of the correlation. ATAC-seq tracks were generated from pseudo-bulk chromatin profiles of cardiomyocytes 1 and cardiomyocytes 2. H3K3me1 ChIP-seq track was generated from an adult non-failing heart (Gilsbach, Ralf et al. 2018). Binding sites of NEF2L1 supported by ATAC-seq footprints are highlighted. **d-e.** Same as in **c.** but for COL1A1 and CREB3L2. Binding sites of RUNX2 supported by ATAC-Seq footprints highlighted. **f.-g.** MISTy's mean importances of the spatial expression (paraview) of ECM genes and cytokines to predict the expression of marker genes across different cardiomyocytes and fibroblast subtypes.

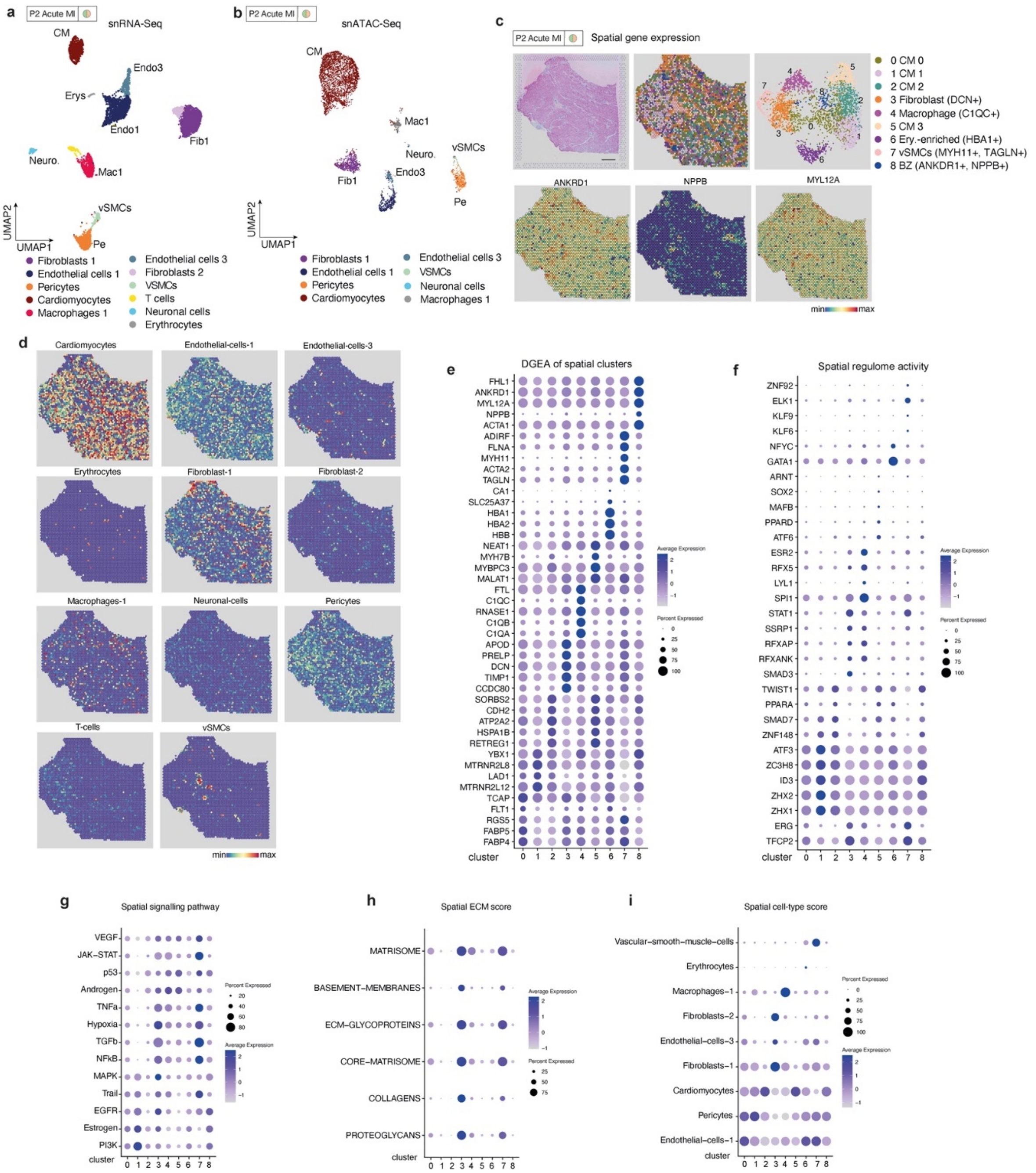


**Extended Data Fig. 11 Spatial characterization of acute myocardial infarction sample border zone.** **a.** UMAP embedding of the snRNA-Seq and **b.** snATAC-seq data. **c.** Clustering of the spatial spots resulted in 9 clusters (upper left and middle panel). UMAP embedding of the slide spots (upper right panel). Spatial distribution of the expression of ANKRD1, NPPB, and MYL12A across the detection area. **d.** Cell-type score per spot by label transfer. Coverage and mean expression (activity or score) per cluster in spatial transcriptomics of: **e.** The top 5 differentially expressed genes of each cluster, **f.** The top 5 differentially active TFs using DoRothEA, **g.** PROGENy's signaling pathway activities, **h.** ECM scores, and **i.** Cell-type scores as calculated with label transfer (see Methods).

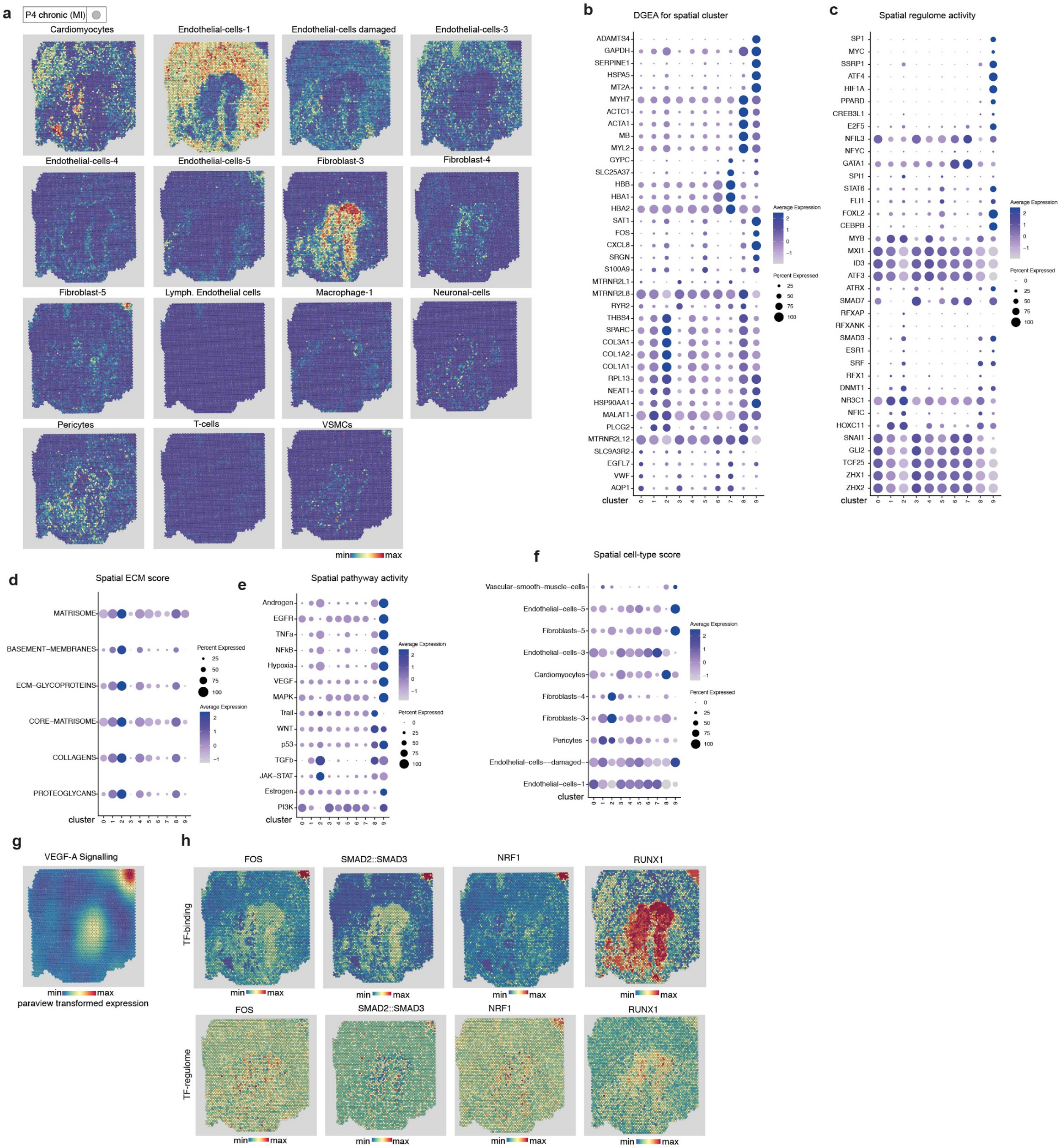


**Extended Data Fig. 12 Characterization of chronic fibrotic zones with temporal scar differences. a.** Cell-type score per spot by label transfer. Coverage and mean expression (activity or score) per cluster in spatial transcriptomics of: **b.** The top 5 differentially expressed genes of each cluster, **c.** The top 5 differentially active TFs using DoRothEA, **d.** ECM scores, **e.** PROGENy's signaling pathway activities, and **f.** Cell-type scores as calculated with label transfer (See Methods). **g.** Paraview transformed spatial activities (l=10, see Methods) of PROGENy's VEGF pathway. **h.** Spatial HINT TF-Binding activity (upper) and summarized gene expression of target genes (lower) using module scores of FOS, SMAD2/SMAD3, NRF1, and RUNX1.

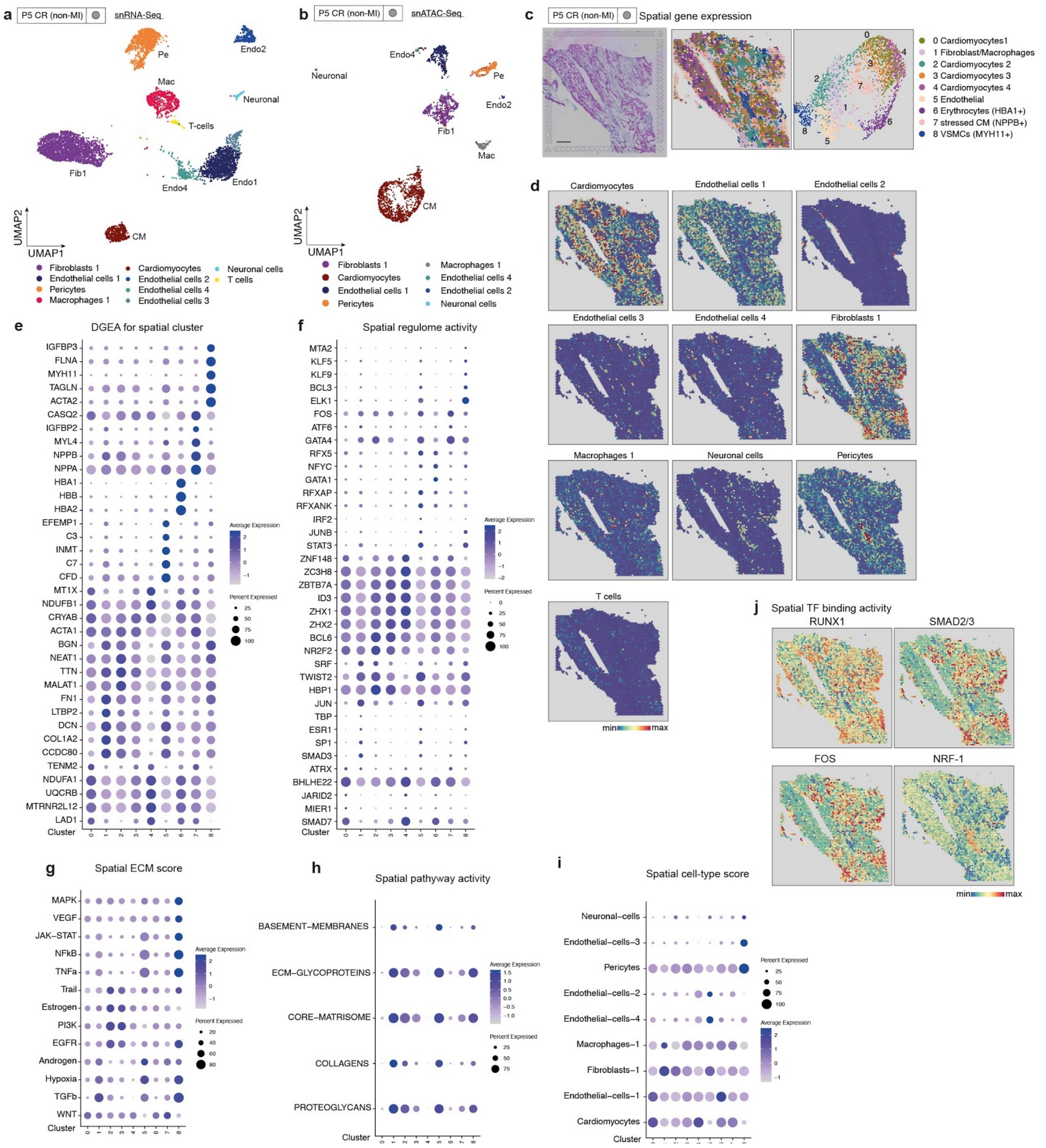


**Extended Data Fig. 13. Characterization of chronic fibrotic zones with temporal scar differences in non-MI heart tissue.** **a.** UMAP embedding of the snRNA-Seq and **b.** snATAC-seq data. **c.** Clustering of the spatial spots resulted in 9 clusters (middle panel). UMAP embedding of the slide spots (upper right panel). **d.** Cell-Type score per spot by label transfer. Coverage and mean expression (activity or score) per cluster in spatial transcriptomics of: **e.** The top 5 differentially expressed genes of each cluster, **f.** The top 5 differentially active TFs using DoRothEA. **g.** PROGENy's signaling pathway activities, **h.** ECM scores, and **i.** Cell-type scores as calculated with label transfer (see Methods).

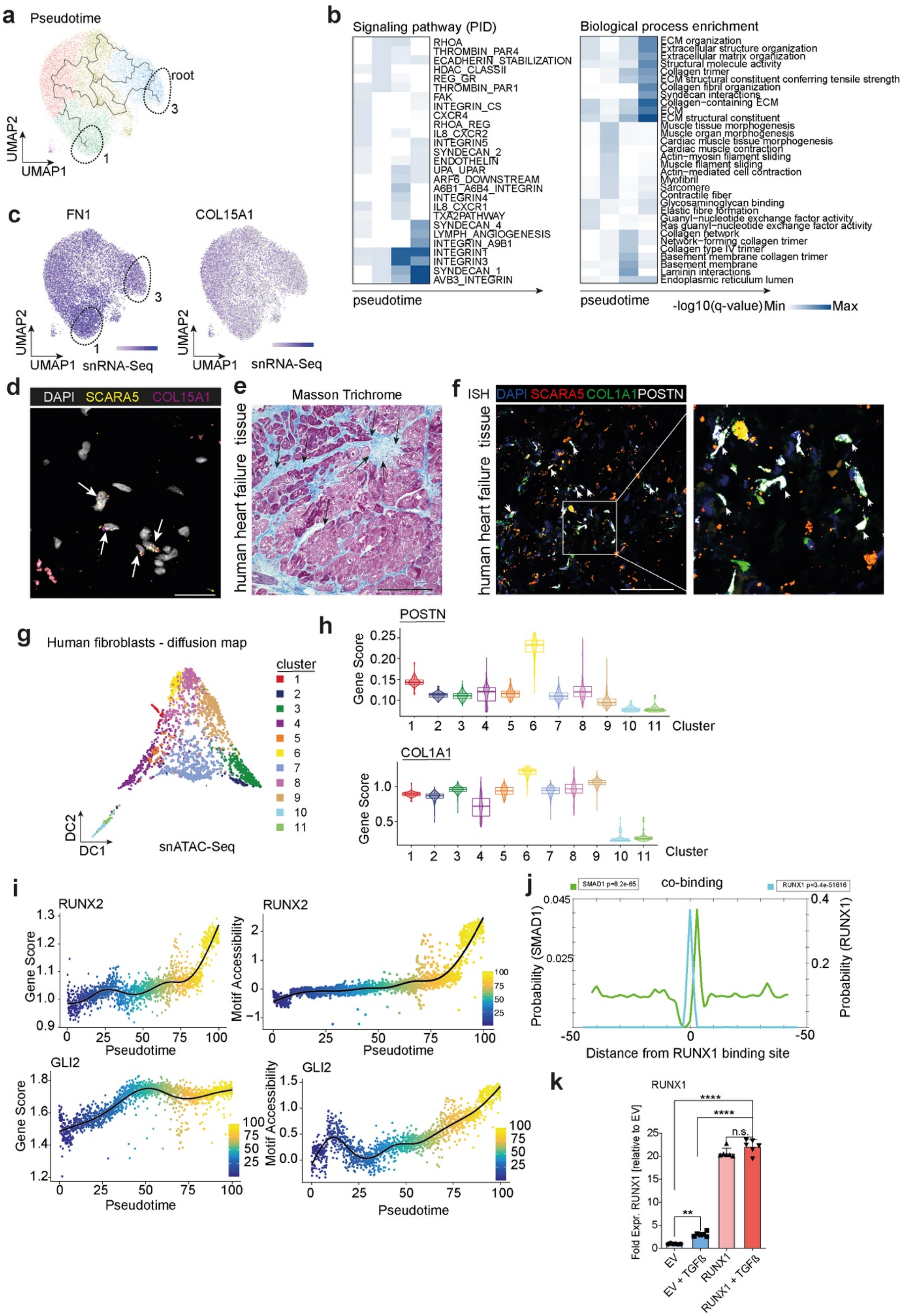

a Pseudotime
root
UMAP2
UMAP1
b Signaling pathway (PID)
RHOA
THROMBIN_PAR4
ECADHERIN_STABILIZATION
HDAC_CLASSII
REG_GR
THROMBIN_PAR1
FAK
INTEGRIN_CS
CXCR4
RHOA_REG
IL8_CXCR2
INTEGRIN5
SYNDECAN_2
ENDOTHELIN
UPA_UPAR
ARF6_DOWNSTREAM
A6B1_A6B4_INTEGRIN
INTEGRIN4
IL8_CXCR1
TXA2PATHWAY
SYNDECAN_4
LYMPH_ANGIOGENESIS
INTEGRIN_A9B1
INTEGRIN1
INTEGRIN3
SYNDECAN_1
AVB3_INTEGRIN
pseudotime
Biological process enrichment
ECM organization
Extracellular structure organization
Extracellular matrix organization
Structural molecule activity
Collagen trimer
ECM structural constituent conferring tensile strength
Collagen fibril organization
Syndecan interactions
Collagen-containing ECM
ECM
ECM structural constituent
Muscle tissue morphogenesis
Muscle organ morphogenesis
Cardiac muscle tissue morphogenesis
Cardiac muscle contraction
Actin-myosin filament sliding
Muscle filament sliding
Actin-mediated cell contraction
Myofibril
Sarcomere
Contractile fiber
Glycosaminoglycan binding
Elastic fibre formation
Guanyl-nucleotide exchange factor activity
Ras guanyl-nucleotide exchange factor activity
Collagen network
Network-forming collagen trimer
Collagen type IV trimer
Basement membrane collagen trimer
Basement membrane
Laminin interactions
Endoplasmic reticulum lumen
pseudotime
-log10(q-value) Min Max
c FN1
COL15A1
UMAP2
UMAP1 snRNA-Seq
d DAPI SCARA5 COL15A1
e Masson Trichrome
human heart failure tissue
f ISH DAPI SCARA5 COL1A1 POSTN
g Human fibroblasts - diffusion map
cluster
DC2
DC1
snATAC-Seq
h POSTN
Gene Score
Cluster
COL1A1
i RUNX2
Gene Score
Pseudotime
Motif Accessibility
GLI2
j co-binding
SMAD1 p=8.2e-65
RUNX1 p=3.4e-51616
Probability (SMAD1)
Probability (RUNX1)
Distance from RUNX1 binding site
k RUNX1
Fold Expr. RUNX1 [relative to EV]
EV
EV + TGFß
RUNX1
RUNX1 + TGFß
n.s.

**Extended Data Fig. 14 Multi-omics trajectory analysis of myofibroblast differentiation. a.** UMAP showing sub-clusters and pseudotime of the integrated fibroblasts from snRNA-seq data. **b.** Scatter plot showing gene expression of FN1 (left) and COL15A1 (right). **c.** Pathway enrichment across pseudotime for lineage in a.. Biological process enrichment along pseudotime for lineage in a. **d.** Representative in-situ hybridisation of SCARA5 and COL15A1 on human heart tissue. Scale bars: 30 µm. **e.** Representative Masson trichrome staining of a human heart sample stained in f. Clinical data in Supplementary Table 2**.** Scale bar = 50 µm. **f.** Representative in-situ of human heart failure tissue showing SCARA5, COL1A1 and POSTN expression. Scale bar = 50 µm. **f.** Quantification and comparison of POSTN+/SCARA5+ cells vs. POSTN+/COL1A1+ cells in human heart failure tissues (n=7). Mann-Whitney test. **g.** Diffusion map showing sub-clusters of the integrated fibroblasts from snATAC-seq data. **h.** Violin plot showing gene score of POSTN and COL15A1 across all sub-clusters. **i.** Gene score (left) and motif activity (right) along the trajectory of GLI2 (upper) and RUNX2 (lower). Each dot represents an individual pseudotime-ordered cell. **j.** Line plot showing binding probability SMAD1 and RUNX1. Genomic coordinates were obtained using RUNX1 binding sites and sequences were extracted from the reference genome hg38. The signal was smoothed based on a window size (n = 5). **k.** Expression of RUNX1 by qPCR after RUNX1 overexpression with and without TGFβ compares to empty vector (EV). n=6. One-way ANOVA followed by Bonferroni's correction.

**Supplementary File 1**: Differential analysis of gene expression, transcription factor regulome-based activities (DoRothEA), pathway activities, and cell-type and ECM scores of each cluster of each spatial transcriptomics dataset.

**Supplementary File 2**: Marker List for MISTy

**Supplementary File 3**: Cell-type-specific TF binding activity

**Extended Data Table 1**: Primer qPCR

| **Genes** | **Forward Primer** | **Reverse Primer** |
|---|---|---|
| *GAPDH* | 5‘-GAAGGTGAAGGTCGGAGTCA | 5‘-TGGACTCCACGACGTACTCA |
| *collagen type I alpha 1 chain* | 5‘-CCCAGCCACAAAGAGTCTACA | 5‘-ATTGGTGGGATGTCTTCGTCT |
| *fibronectin 1* | 5‘-AACAAACACTAATGTTAATTGCCCA | 5‘-TCGGGAATCTTCTCTGTCAGC |
| *actin alpha 2, smooth muscle* | 5‘-ACTGCCTTGGTGTGTGACAA | 5‘-CACCATCACCCCCTGATGTC |
| *RUNX1* | 5‘-CAGTCGACTCTCAACGGCAC | 5‘-TAGGTGAAGGCGCCTGGATA |

**Extended Data Table 2**: Clinical data heart failure patients

| Patient No. | Sex | Age (in years) | Diagnosis | LV-EF in % | Masson-Trichrome stained area in % |
|---|---|---|---|---|---|
| 1 | Male | 74 | ICM | 15% | 19.91 |
| 2 | Male | 53 | ICM | 18% | 12.23 |
| 3 | Male | 50 | DCM | 16% | 10.34 |
| 4 | Male | 45 | ICM | 19% | 8.19 |
| 5 | Male | 67 | ICM | 31% | 6.97 |
| 6 | Male | 65 | ICM | 12% | 16.35 |
| 7 | Male | 68 | ICM | 25% | 8.3 |

ICM= Ischemic cardiomyopathy, DCM= dilatative cardiomyopathy, LV-EF= Left ventricular ejection fraction in %

### Material and Methods

#### Ethics

The local ethics committee of the Ruhr University Bochum in Bad Oeynhausen and of the RWTH Aachen University approved all human tissue protocols (No 220-640, EK151/09 respectively). Human myocardial tissue was collected from non-transplanted donor hearts or patients after myocardial infarction undergoing heart transplantation or implantation of a total artificial heart. All patients provided informed consent and the study was performed in accordance with the Declaration of Helsinki.

#### Human tissue processing and screening

Heart tissues were sampled by the surgeon and immediately frozen in liquid nitrogen. Tissues were dounced in liquid nitrogen and 7-$10^3$ pieces were embedded in O.C.T. compound (Tissue-Tek) and frozen on dry-ice. 10 µm cryo tissue sections were HE stained and the appropriate tissue regions selected for further processing. 25 human tissue samples were screened. For RNA quality control we minced a 3x3 $mm^3$ heart tissue piece in liquid nitrogen and isolated the RNA using Qiagen RNeasy Mini kit (Qiagen) using a proteinase K digestion step as suggested in RNeasy Fibrous Tissue Mini Kit (Qiagen, 74704). RNA Integrity Number analysis (Agilent) was performed using Bioanalyzer RNA 6000 Nano kits (Agilent, No. 5067).

#### Spatial gene expression assay 10X Genomics Visium

Frozen heart samples were embedded in OCT (TissueTek) and cryosectioned (Thermo Cryostar). The 10 µm section were placed on the prechilled Optimization slides and reagent kit (Visium, 10X Genomics, PN-1000193) and after the optimization procedure on the Spatial Gene expression slides (Visium, 10X Genomics, PN-1000187). The tissues were treated as recommended by the 10X Genomics and the optimization procedure showed an optimal permeabilization time of 12 or 18 minutes of digestion and release of RNA from the tissue slide. Brightfield histological images were taken using a 10X objective on the Nikon Eclipse TiE. Stitching of the raw images was performed using the NIS-Elements software. NGS-libraries were prepared according to the Visium User Guide. Libraries were loaded at 300 pM and sequenced on a NovaSeq 6000 System (Illumina) as recommended by 10X Genomics.

#### Single nuclei isolation of human hearts and FANS

Single nuclei isolation was performed as previously described[1]. In brief, heart tissue was cut into small pieces (0,5 $mm^3$) in a sterile petri dish on ice and transferred to a tissue douncer. Nuclei isolation buffer 0,5 ml (EZ lysis buffer, NUC101, Sigma-Aldrich) plus RNAse inhibitor (Protector RNase Inhibitor, Roche) were added to the tissue and 10-15 strokes with pestle A were applied followed by 10-15 strokes of pestle B. The nuclei were stained with DAPI and FACS sorted using SONY SH800 to enrich nuclei.

#### Single cell assay 10X Genomics 3' sc-RNA-seq

Nuclei suspensions with a concentration ranging from 400-1000 nuclei/ul were loaded into the chromium controller (10X, Genomics, PN-120223') on a Single cell B chip (10X Genomics, PN-120262) and processed following the manufacturer' original protocol to generate GEMs (single cell gel beads in emulsion). The sequencing library was generated using the Chromium Single cell 3' reagent Kit v3 (10X, PN-1000092) and Chromium i7 Multiplex Kit (10X Genomics, PN-120262). Quality control for the constructed library was performed by Tape station (company). Libraries were sequenced on NovaSeq targeting 50k reads per cells 2x150 paired-end kits using the following read length: 28 bp Read1 for cell barcode and UMI, 8 bp I7 index for sample index and 91 bp Read2 for transcript.

**Single cell assay 10X Genomics 3' sc-ATAC-seq**

The remaining nuclei after processing for 3' sc-RNA-Seq Assay were centrifuged 500g 4°C for 5 min and resuspended in 10 ul of nuclei suspension buffer. After tagmentation the nuclei suspension was loaded on the Chromium Chip E (10X Genomics, PN-1000082) in the Chromium Controller according to the manufacturer's protocol. The library was sequenced on an Illumina Novaseq 2x50 paired-end kits using the following read length: 50 bp Read1 for DNA fragments, 8 bp for i7 index for sample index, 16 bp i5 index for cell barcodes and 50 bp Read2 for DNA fragments.

**RNA in-situ hybridization and image quantification**

In situ hybridization was performed using formalin-fixed paraffin embedded tissue samples and the RNAScope Multiplex Detection KIT V2 (RNAScope, #323100) following the manufacturer's protocol with minor modifications. The antigen retrieval was performed for 22 min at 96°C instead of 15 min at 99°C in a water bath. 3-5 drops of pre-treatment 1 solution were incubated at room temperature (RT) for 10 minutes after performing antigen retrieval. The washing steps were performed 5 minutes three times. The following probes were used for the RNAscope assay: Hs-Col1a1 #401891, Hs-Col1a1 #401891-C2, Hs-Postn #409181-C3, Hs-Pecam1 #487381-C2, Hs-Col15a1 #484001-C2, Hs-Scara5 #574781-C3.

**Masson Trichrome staining and quantification**

Masson's trichrome staining was conducted using ready–to-use kit (Trichrome Stain (Masson) Kit, HT15, Sigma-Aldrich) as described by the manufacturer. Color-based thresholding of blue regions in ImageJ[2] was used to determine the Integrated Density of the collagen positive tissue regions.

**Antibodies and immunofluorescence stainings**

Heart tissues were fixed in 4% formalin for 4 hours at RT and then embedded in paraffin. For staining slides were blocked in 5% donkey serum followed by 1-hour incubation of the primary antibody, washing 3 times for 5 minutes in PBS and subsequent incubation of the secondary antibodies for 45 minutes. Following DAPI (4′,6′-′diamidino-2-phenylindole) staining (Roche, 1:10.000) the slides were mounted with ProLong Gold (Invitrogen, #P10144). The following antibodies were used: anti-ACTA2(aSMA)-Cy3 (C6198,1:250, Sigma-Aldrich), anti-SEMA3G (HPA001761, 1:100, Sigma Aldrich), AF647 donkey anti-rabbit (1:200, Jackson Immuno Research).

**Confocal imaging**

Acquisition of images was performed using a Nikon A1R confocal microscope using 40x and 60x objectives (Nikon). Image processing was performed using the Nikon Software or ImageJ[2].

**Generation of a human PDGFRb$^+$ cardiac cell-line**

PDGFRb$^+$ cells were isolated from a 69 years old male patient, undergoing left ventricular assist device surgery. To generate a single cell suspension, the tissue was homogenised in a gentleMACS dissociator (Miltenyi) and digested with liberase (200 µg/mL, Roche #5401020001) and DNase (60 U/mL), for 30 minutes at 37°C. After filtering the cell suspension (70 µm mesh), cells were stained in two steps using a specific PDGFRb antibody (R&D # MAB1263 antibody, dilution 1:100) followed by Anti-Mouse IgG1-

MicroBeads solution (Miltenyi, #130-047-102). Following MACS, cells were cultured in DMEM media (Thermo Fisher # 31885) for 20 days and immortalized using SV40LT and HTERT. Retroviral particles were produced by transient transfection of HEK293T cells using TransIT-LT (Mirus). Two types of amphotropic particles were generated by co-transfection of plasmids pBABE-puro-SV40-LT (Addgene #13970) or xlox-dNGFR-TERT (Addgene #69805) in combination with a packaging plasmid pUMVC (Addgene #8449) and a pseudotyping plasmid pMD2.G (Addgene #12259). Retroviral particles were 100x concentrated using Retro-X concentrator (Clontech) 48hrs post-transfection. Cell transduction was performed by incubating the target cells with serial dilutions of the retroviral supernatants (1:1 mix of concentrated particles containing SV40-LT or rather hTERT) for 48hrs. Subsequently the infected PDGFRb+ cells were selected with 2 µg/ml puromycin at 72 h after transfection for 7 d.

**Lentiviral overexpression of Runx1**

The human cDNA of *RUNX1* was PCR amplified using the primer sequences 5'- atgcgtatccccgtagatgcc -3' and 5'- tcagtagggcctccacacgg -3'. Restriction sites and N-terminal 1xHA-Tag have been introduced into the PCR product using the primer 5'- cactcgaggccaccatgtacccatacgatgttccagattacgctcgtatccccgtagatgcc -3' and 5'- acggaattctcagtagggcctccacac -3'. Subsequently, the PCR product was digested with XhoI and EcoRI and cloned into pMIG (pMIG was a gift from William Hahn (Addgene plasmid # 9044 ;http://n2t.net/addgene:9044 ; RRID:Addgene_9044). Retroviral particles were produced by transient transfection in combination with packaging plasmid pUMVC (pUMVC gifted from Bob Weinberg (Addgene plasmid # 8449)) and pseudotyping plasmid pMD2.G (pMD2.G gifted from Didier Trono (Addgene plasmid # 12259 ; http://n2t.net/addgene:12259 ; RRID:Addgene_12259)) using TransIT-LT (Mirus). Viral supernatants were collected 48-72 hours after transfection, clarified by centrifugation, supplemented with 10% FCS and Polybrene (Sigma-Aldrich, final concentration of 8µg/ml) and 0.45µm filtered (Millipore; SLHP033RS). Cell transduction was performed by incubating the PDGFß cells with viral supernatants for 48hrs. eGFP-expressing cells were single-cell sorted.

**Quantitative RT-PCR**

Cell pellets were harvested and washed with PBS followed by RNA extraction according to the manufacturer's instructions using the RNeasy Mini Kit (Qiagen). 200 ng total RNA was reverse transcribed with High-Capacity cDNA Reverse Transcription Kit (Applied Biosystems) and qRT-PCR was carried out further as previously described[3] Data were analyzed using the 2-CT method. The primers used are listed in Extended Data table 1.

**snRNA-seq data processing**

Raw snRNA-seq FASTQ files were aligned to the human genome GRCh38 and feature counts were generated with *Cell Ranger* version 3.1.0 with default settings (10X Genomics). To account for the presence of unspliced pre-mRNA in snRNA-seq protocols, a custom "pre-mRNA" reference package was built where reads mapping to both introns and exons were counted (https://support.10xgenomics.com/single-cell-gene-expression/software/pipelines/latest/advanced/references). Downstream analysis and quality control was conducted on the filtered feature-barcode matrices using *Seurat* version 3.1.5 [4]. Nuclei with < 200 or > 2500 detected genes were excluded from the analysis. Nuclei with < 300 unique molecular identifiers (UMIs) and mitochondrial gene mapping > 5% were also filtered out. For every dataset, each feature was log-normalized and scaled with default settings.

**Clustering and cell annotation of snRNA-seq data**

To provide markers for the cell-type label transfer to spatial transcriptomics data, the snRNA-seq datasets were clustered and annotated separately. Dimensionality reduction was performed on the 2,000 most variable features with a principal component analysis (PCA). A shared nearest neighbor (SNN) graph was built with the first 20 principal components (PCs) using *FindNeighbors*, and the cells were clustered with a Louvain-based algorithm with *FindClusters*. The clusters were embedded and visualized in a Uniform Manifold Approximation and Projection (UMAP) using the top 20 PCs[5]. Based on the number of UMIs and detected features, low quality clusters were removed, and the IZ snRNA-seq sample of patient 3 *P3 IZ* was excluded from the analysis. Manuel high level cell annotation of the retained clusters was based on canonical cell type markers[6–8] (Extended data Fig. 4a). To characterise cell-subtypes, each gene was ranked in every cluster based on uniqueness and level of expression (specificity score) by *sortGenes* from the *genesorteR* package[9]. The top 50 ranking genes for each cluster with a conditional probability of expression > 0.35 were considered for the cell annotations.

**Differential gene expression between cardiomyocytes subpopulations in the border zone**

Differentially expressed genes between cardiomyocytes 1 and 2 in the snRNA-seq data of the border zone (P3_BZ) were estimated with *FindMarkers* (min.pct = 0.25). Significant genes (FDR < 0.05) were visualized in a volcano plot and important genes were highlighted (Extended data Fig. 9e).

**snRNA-seq data integration**

To ensure a uniform cell annotation of the individual snRNA-seq datasets, the samples were integrated. The 2,000 most variable features were used to determine shared correlation structures through a canonical correlation analysis using *FindIntegrationAnchors* with 20 dimensions [10]. The anchoring features were used to integrate the cells by *IntegrateData* using 20 dimensions. The data was scaled and visualized in a UMAP based on the first 20 PCs. The robustness of the cell annotations from the individual datasets was evaluated by calculating the Pearson correlation coefficients of the average gene expression of the integrated data (Fig. S3A).

**Trajectory analysis of sub-clustered fibroblasts**

In order to study the differentiation process of myofibroblasts in the snRNA-seq data, the fibroblast subpopulations of the integrated data were further sub-clustered following the previously described protocol in Seurat. Pseudotime analysis was performed with Monocle 3[11] on the UMAP embedding of the sub-clustered fibroblasts. Based on our previous work[12], cells with the highest expression of *SCARA5* were chosen as the root in the trajectory and the cells were ordered in pseudotime accordingly. Differentially expressed genes along the trajectory were determined with the Moran's I test via the function *graph_test* (neighbor_graph = "principal_graph"). The top 40 genes, ordered by the Moran's test statistic (q < 0.05), were clustered with the *k*-means algorithm and visualized along pseudotime with the *genesorteR* function *plotMarkerHeat* (Fig. 6d).

**Pathway and gene ontology enrichment analysis**

Pathway and gene ontology (GO) enrichment analysis was performed on the sub-clustered fibroblast. The cells were ordered based on pseudotime and were partitioned into 4 equal-sized bins where *FindAllMarkers* (min.pct = 0.25) was used to determine differentially expressed genes. Significant genes (FDR < 0.05) were ordered by log2FC and GO enrichment analysis was performed on each bin with the *gost* function from the *gprofiler2* package[13]. Electronic annotations were excluded from the analysis. For the pathway analysis, PID data was downloaded from MSigDB[14] in November 2020 as “gmt” files and the pathway enrichment for each bin was likewise computed with *gost*. The top GO and pathway terms for each bin were visualized with *heatmap.2* (Extended data Fig. 14c).

**ECM scores**

ECM and collagen scores were computed with Seurat’s *AddModuleScore* as the average expression levels of core matrisome and collagen genes provided in [15] subtracted by the aggregated expression of 35 randomly selected control genes[16].

**snATAC-seq data processing**

Low level processing of snATAC-seq data, including demultiplexing, reads trimming, filtering, alignment, barcode counting and peak calling, was performed using the *cellranger atac* pipeline (version 1.1.0) with default setting for each sample. We used the fragments and initial cell detection in the following analysis and excluded sample *P3 IZ* due to low quality of the snRNA-seq data.

**Estimation of gene activity using snATAC-seq and alignment with snRNA-seq**

A gene activity matrix was created for each sample by summing the reads intersecting the gene body and promoter region (2kb upstream). The gene coordinates were extracted from EnsDb.Hsapiens.v86[17] and the fragments file was used as input to count the number of fragments that map to each of these regions for each cell using the *FeatureMatrix* function from Signac version 0.2.5[18]. To obtain a consistent identification of cell populations, the snRNA-seq data from the same sample was used to annotate the snATAC-seq data using the label transfer approach of Seurat[4]. First, a Seurat object was created using the function *CreateSeuratObject* (min.cell = 1) and only cells with a number of unique fragments > 3,000 were kept for analysis. Then, term frequency inverse document frequency (TF-IDF) normalization was performed using function *RunTFIDF* from Signac and dimensionality reduction was done with function *RunSVD*. The first component was highly correlated with the number of fragments and was therefore removed from our downstream analysis. Next, a set of anchors between snRNA- and snATAC-seq were identified using the function *FindTransferAnchors* and the annotated cell-type labels from snRNA-seq were used as reference for label transferring using the function *TransferData*.

**snATAC-seq re-clustering and re-annotation**

To better identify rare cell-types and increase the power of clustering analysis of snATAC-seq, a second round of clustering and label transferring was performed. Specifically, the predicted cell labels from the first round were used to group the cells, and for each group, fragments were extracted and peak calling was performed with MACS2[19]. The most significant 100,000 summits were selected and extended to 500 bp (±250). To create a union peak set, all selected summits were iteratively merged and the overlapping peaks were removed using the approach described in [20]. The ENCODE blacklist of hg38 was used to further filter the peaks[21]. A count matrix was created using the union peak set as feature and fragments as input. The new matrix contained more specific regulatory regions than the matrix generated by the Cell Ranger pipeline and gene activity matrix. This was used for the second round of cell clustering and label transferring as previously described. Finally, each cluster was annotated by using the most representative cell-type predicted by the label transfer approach. To visualize the snATAC- and snRNA-seq data in a common low dimensional space, the anchors used in the second round to transfer labels were used to impute the gene activity in snATAC-seq. This imputed gene activity was merged to snRNA-seq and a standard Seurat analysis was performed to visualize all cells together in a co-embedding.

**Identification of peak-to-gene links**

We used a procedure from [20] to associate peaks to genes, which is based on first finding neighboring snRNA- and snATAC-seq in a common embedding and identifying peak-to-gene pairs with high correlation between their expression and accessibility. For this, we only considered gene-peak pairs within +/-250kbp by using the function *findOverlaps* from GenomicsRangers[22]. For each cell from snATAC-seq data, the nearest neighbor from snRNA-seq data was identified by using the co-embedding as input. Due to the low coverage nature of reads in snATAC-seq cells, we used the sum of 50 nearest snATAC-seq cells. The two matrices (snATAC-seq aggregate matrix and snRNA-seq neighbors matrix) shared the same number of cells, allowing the calculation of the correlation between genes (snRNA-seq) and peaks (ATAC-seq aggregate). To test if the correlation was significant, for each gene, 1,000 peaks from a different chromosome were selected and a null model was created. Under the assumption that the correlation was normally distributed, a z-score test was performed to compute a p-value. Only significant links ($FDR < 0.05$) with positive correlation and distance to a gene promoter > 2,000 bp were kept. *ComplexHeatmap* was used to visualize all predicted links for each sample. The columns were first ordered by cell-type, followed by a clustering for each cell-type using the function *hclust* with 1 - correlation as distance. Rows were ordered by clustering partitioning around medoids (PAM) algorithm. Number of medoids was set as the number of cell-types identified by snATAC-seq and 1 - correlation was used as distance.

**Differential footprinting analysis and TF regulome prediction**

Differential footprinting was performed using HINT-ATAC[23] to compare TF accessibility scores between distinct cell-types of a snATAC-seq sample. For each cell-type, the reads were extracted to compile a cell-type-specific BAM file using a custom python script. Next, peaks were detected using MACS2[19] and footprints were predicted using HINT-ATAC, followed by motif matching to identify TF binding sites (TFBSs) of all motifs from JASPAR[24]. The TF accessibility score, a measurement of strength of TF binding in a particular biological condition, was calculated using the BAM files and predicted TFBSs. We only considered motifs with at least 1,000 binding sites.

Cell-type-specific TF binding sites were associated with nearby genes using the function *gene_association* from RGT and hg38 as reference genome. The maximum distance between TF binding sites and an associated gene was set to 100kbp. This allowed us to generate cell- and TF-specific regulomes.

**snATAC-seq data integration**

To integrate all snATAC-seq libraries, the fragments of valid cells from all samples were collected to form an aggregated dataset using the *cellranger-atac aggr* command and peaks were detected again based on the aggregated data. A Seurat object was created and for each sample, the data was normalized using the function *NormalizeData* with "LogNormalize". The 5,000 most variable features were identified using the function *FindVariableFeatures* and used to find anchors and integrate the data across all samples. PCA was performed using the function *RunPCA* (npcs = 30) with the integrated data as input. For visualization, the function *RunUMAP* (metric = “correlation”, min.dist = 0.2) was used.

**Processing of cardiac myocytes H3K4me1 ChIP-seq data**

Raw data of cardiac myocytes H3K4me1 ChIP-seq[25] was downloaded from the National Center for Biotechnology Information under the BioProject ID PRJNA353755. Adapter sequences were trimmed from FASTQ files and reads were aligned to hg38 using Bowtie2[26]. All reads mapped to chrY, mitochondria and unassembled random contigs were removed. Duplicates were removed using Picard (https://broadinstitute.github.io/picard/). Reads were further filtered by map quality > 30 and to be properly paired. Finally, a bigwig file was generated using *deepTools*[27] for visualization (normalizeUsing CPM).

**Sub-clustering and trajectory analysis of fibroblast snATAC-seq data**

Sub-clustering and trajectory analysis were performed using ArchR[28] (https://www.archrproject.com/). The fragments of fibroblasts from all samples were collected and an ArchR project was created. Dimensionality reduction was performed using the function *addIterativeLSI* and batch effects were corrected using the function *addHarmony*. To identify a trajectory, a diffusion map was generated using the package *destiny*[29]. Sub-clusters and marker genes were identified via the function *addClusters* and *getMarkerFeatures*. Peaks were detected for each sub-cluster using the function *addReproduciblePeakSet* and cell-specific TF motif deviation scores were inferred using ChromVAR[30]. The cellular trajectory was created from $SCARA5^+$ to $POSTN^+$ cells using the function *addTrajectory*, and heatmaps of gene scores and motif deviation scores along the trajectory were plotted using the function *plotTrajectoryHeatmap*. Finally, an integrative analysis for identification of positive regulators was performed using the function *correlateTrajectories* and TFs with significant correlation (FDR < 0.1) were shown.

**RUNX1 and SMAD1 co-binding analysis**

To test if RUNX1 was co-binding with SMAD1 in myofibroblasts, we collected all RUNX1-binding sites from myofibroblasts and extended 50bps for both sides. We performed local motif enrichment analysis using *CentriMo*[31] and used RUNX1 and SMAD1 motifs as input. The enrichment p-value was calculated using a one-tailed binomial test and was corrected for multiple tests using Bonferroni correction.

**Characterization of spatial transcriptomics data sets**

Filtered expression matrices from *Space Ranger* were used as initial input for the spatial transcriptomics analysis. Individual count matrices were normalized with *sctransform* implemented in Seurat 3.1.4.9[4]. For each spot, we estimated signaling pathway activities with PROGENy's model matrix using the top 1,000 genes of each transcriptional footprint[32,33]. Additionally, TF activities were estimated with *viper*[34] for regulons obtained from DoRothEA[35]. ECM scores were calculated for each spot as described previously. As a complementary analysis, for each slide, genes with spatial expression patterns were obtained with SPARK[36] (FDR < 0.05). Overrepresented canonical pathways in each set of genes were obtained with hypergeometric tests (FDR < 0.05). Canonical pathways were downloaded from MSigDB[14] in December 2019.

To find groups of spots with similar expression patterns, PCA was applied to the expression matrices. An SNN graph between spots was obtained with the first 30 PCs and clusters were defined with a resolution value of 1 with a Louvain-based algorithm.

Differentially expressed genes, active TFs and signaling pathways, and ECM scores were obtained for each cluster using Wilcoxon tests (FDR < 0.005). In supplemental figures, only the top 5 differentially expressed genes and active TFs are shown (full results available in Supplemental file 1). Mitochondrial genes, although tested, were excluded from all visualizations.

**Estimation of cell-type composition**

To assign cell-type scores to every spot of each slide, we leveraged the information captured in the integrated single nuclear data sets. We transferred the cell-type labels from each sample's single nuclear integrated data set to their corresponding visium slide using Seurat's label transfer approach[4]. Wilcoxon tests were used to compare differences of the cell-type scores between clusters. Label transfer cell-type scores for sample *P3 IZ* were not obtained since snRNA-seq data was unavailable.

**Estimation of the impacts of the spatial context in gene expression and cell-type location**

We used MISTy to find the interactions that the broader tissue structure has with selected cell-types or functional areas within a slide that can explain the expression of marker genes. First, for a major cell-type of interest *c* with multiple subtypes, we identified their locations in an individual slide leveraging its label transfer cell-type scores. Then, we identified all gene expression markers identified in the slide's paired snRNA-seq data that were consistent with the cell-type assignment of spots. Finally, we build MISTy pipelines to predict the expression of the selected gene markers using two different spatial contexts: 1) An intrinsic view that measures the relationships between the selected markers within a spot and 2) an optimized paraview that weights the expression of putative ECM proteins, cytokines or pathway activities in the surroundings of each spot.

Predicted gene markers were defined by the union of the top 50 most specific genes (based on *genesorteR* specificity scores) of the tested cell-types from the paired snRNA-seq data. Additional filtering was done to ensure that these predicted markers were part of the most variable genes of the slide. All spots where the maximum cell-type score belonged to one of the predicted cell-types were considered in the model. In the case of the slides where clusters were tested, the union of the top 20 most differentially expressed genes of each cluster were used as predicted gene markers. Predictor cytokines were obtained from MSigDB's[14] KEGG collection "KEGG_CYTOKINE_CYTOKINE_RECEPTOR_INTERACTION". Predictor ECM proteins were obtained from the NABA collection [15]. Predictor pathway activities were estimated with PROGENy as described previously. Initially, ECM and cytokines with a coverage of at least 10% in the slide were used as predictors in a single MISTy pipeline. Complementary, to capture lowly expressed cytokines, another pipeline was built using only cytokines with a coverage of at least 1% in the slide as predictors. Pathways were used as predictors in an independent pipeline.

Gene markers whose prediction was increased with the inclusion of the paraview were recovered (threshold p-value of gain of explained variance of the multiview in contrast to a single, intrinsic view set at 0.25) and grouped based on their subtype of origin. The mean importances of each view were used as a proxy of the influence of each predictor to subtypes. High importances in the intrinsic view capture representative genes of each cell subtype and high importances in the paracrine view capture relationships between the location of a subtype and the presence of ECM proteins, cytokines or pathway activities in the surrounding microenvironment. Only for visualization purposes, paraview-transformed gene expression or pathway activity was fixed with a parameter $l = 10$.

**Footprinting TF binding activity and regulomes in space**

We mapped the cell-type-specific footprinting-based TF binding activity from snATAC-seq to spatial data for each sample. Specifically, for each spot $i$ and TF $j$, we calculated the activity as following $ACT_{ij} = \sum_{k=1}^{K} Score_{ik} \cdot ACT_{kj}'$, where $Score_{ik}$ is the transferred score for cell-type $k$ and $K$ is the number of cell-types identified from snATAC-seq, $ACT_{kj}'$ is the binding activity of TF $j$ in cell-type $k$ from snATAC-seq. Cell- and sample-specific footprinting-based regulons (see Differential footprinting analysis and TF regulome prediction) were used to validate spatial TF activity in both snRNA and spatial transcriptomics. For this, we estimated regulon scores using the collection of cell-type and sample-specific regulons with viper for the corresponding snRNA-seq and spatial transcriptomics samples. For snRNA-seq, we tested if the cell-type with the highest TF binding activity also showed the highest regulon activity. For spatial transcriptomics, we tested if the top 10% spots with the highest TF binding activity also showed the highest regulon score using Wilcoxon tests.

**References**


1. Habib, N. *et al.* Massively parallel single-nucleus RNA-seq with DroNc-seq. *Nat. Methods* **14**, 955–958 (2017).
2. Schneider, C. A., Rasband, W. S. & Eliceiri, K. W. NIH Image to ImageJ: 25 years of image analysis. *Nat. Methods* **9**, 671–675 (2012).
3. Kramann, R. *et al.* Perivascular Gli1+ progenitors are key contributors to injury-induced organ fibrosis. *Cell Stem Cell* **16**, 51–66 (2015).
4. Comprehensive Integration of Single-Cell Data. *Cell* **177**, 1888–1902.e21 (2019).
5. McInnes, L., Healy, J., Saul, N. & Großberger, L. UMAP: Uniform Manifold Approximation and Projection. *JOSS* **3**, 861 (2018).
6. Litviňuková, M. *et al.* Cells of the adult human heart. *Nature* (2020) doi:10.1038/s41586-020-2797-4.
7. Wang, L. *et al.* Single-cell reconstruction of the adult human heart during heart failure and recovery reveals the cellular landscape underlying cardiac function. *Nat. Cell Biol.* **22**, 108–119 (2020).
8. Tucker, N. R. *et al.* Transcriptional and Cellular Diversity of the Human Heart. *Circulation* (2020) doi:10.1161/CIRCULATIONAHA.119.045401.
9. Ibrahim, M. M. & Kramann, R. genesorteR: Feature Ranking in Clustered Single Cell Data. *Cold Spring Harbor Laboratory* 676379 (2019) doi:10.1101/676379.
10. Butler, A., Hoffman, P., Smibert, P., Papalexi, E. & Satija, R. Integrating single-cell transcriptomic data across different conditions, technologies, and species. *Nat. Biotechnol.* **36**, 411–420 (2018).
11. Cao, J. *et al.* The single-cell transcriptional landscape of mammalian organogenesis. *Nature* **566**, 496–502 (2019).
12. Kuppe, C. *et al.* Decoding myofibroblast origins in human kidney fibrosis. *Nature* (2020) doi:10.1038/s41586-020-2941-1.
13. Raudvere, U. *et al.* g:Profiler: a web server for functional enrichment analysis and conversions of gene lists (2019 update). *Nucleic Acids Res.* **47**, W191–W198 (2019).
14. Liberzon, A. *et al.* The Molecular Signatures Database Hallmark Gene Set Collection. *Cell Systems* vol. 1 417–425 (2015).
15. Naba, A. *et al.* The extracellular matrix: Tools and insights for the 'omics' era. *Matrix Biol.* **49**, 10–

24 (2016).
16. Tirosh, I. *et al.* Dissecting the multicellular ecosystem of metastatic melanoma by single-cell RNA-seq. *Science* **352**, 189–196 (2016).
17. Rainer, J. EnsDb.Hsapiens.v86: Ensembl based annotation package. (2017).
18. Stuart, T., Srivastava, A., Lareau, C. & Satija, R. Multimodal single-cell chromatin analysis with Signac. *Cold Spring Harbor Laboratory* 2020.11.09.373613 (2020) doi:10.1101/2020.11.09.373613.
19. Zhang, Y. *et al.* Model-based analysis of ChIP-Seq (MACS). *Genome Biol.* **9**, R137 (2008).
20. Granja, J. M. *et al.* Single-cell multiomic analysis identifies regulatory programs in mixed-phenotype acute leukemia. *Nat. Biotechnol.* **37**, 1458–1465 (2019).
21. Amemiya, H. M., Kundaje, A. & Boyle, A. P. The ENCODE Blacklist: Identification of Problematic Regions of the Genome. *Sci. Rep.* **9**, 9354 (2019).
22. Lawrence, M. *et al.* Software for computing and annotating genomic ranges. *PLoS Comput. Biol.* **9**, e1003118 (2013).
23. Li, Z. *et al.* Identification of transcription factor binding sites using ATAC-seq. *Genome Biol.* **20**, 45 (2019).
24. Fornes, O. *et al.* JASPAR 2020: update of the open-access database of transcription factor binding profiles. *Nucleic Acids Res.* **48**, D87–D92 (2020).
25. Gilsbach, R. *et al.* Distinct epigenetic programs regulate cardiac myocyte development and disease in the human heart in vivo. *Nat. Commun.* **9**, 391 (2018).
26. Langmead, B. & Salzberg, S. L. Fast gapped-read alignment with Bowtie 2. *Nat. Methods* **9**, 357–359 (2012).
27. Ramírez, F. *et al.* deepTools2: a next generation web server for deep-sequencing data analysis. *Nucleic Acids Res.* **44**, W160–5 (2016).
28. Granja, J. M. *et al.* ArchR: An integrative and scalable software package for single-cell chromatin accessibility analysis. *Cold Spring Harbor Laboratory* 2020.04.28.066498 (2020) doi:10.1101/2020.04.28.066498.
29. Angerer, P. *et al.* destiny: diffusion maps for large-scale single-cell data in R. *Bioinformatics* **32**, 1241–1243 (2016).
30. Schep, A. N., Wu, B., Buenrostro, J. D. & Greenleaf, W. J. chromVAR: inferring transcription-factor-associated accessibility from single-cell epigenomic data. *Nat. Methods* **14**, 975–978 (2017).
31. Bailey, T. L. & Machanick, P. Inferring direct DNA binding from ChIP-seq. *Nucleic Acids Res.* **40**, e128 (2012).
32. Schubert, M. *et al.* Perturbation-response genes reveal signaling footprints in cancer gene expression. *Nat. Commun.* **9**, 20 (2018).
33. Holland, C. H., Szalai, B. & Saez-Rodriguez, J. Transfer of regulatory knowledge from human to mouse for functional genomics analysis. *Biochim. Biophys. Acta Gene Regul. Mech.* **1863**, 194431 (2020).
34. Alvarez, M. J. *et al.* Functional characterization of somatic mutations in cancer using network-based inference of protein activity. *Nat. Genet.* **48**, 838–847 (2016).
35. Garcia-Alonso, L., Holland, C. H., Ibrahim, M. M., Turei, D. & Saez-Rodriguez, J. Benchmark and integration of resources for the estimation of human transcription factor activities. *Genome Res.* **29**, 1363–1375 (2019).
36. Sun, S., Zhu, J. & Zhou, X. Statistical analysis of spatial expression patterns for spatially resolved transcriptomic studies. *Nat. Methods* **17**, 193–200 (2020).
